# Structural brain correlates of poor reading comprehension

**DOI:** 10.1101/2025.07.23.666369

**Authors:** Kelly Mahaffy, Nabin Koirala, Daniel Kleinman, Nicole Landi

## Abstract

Poor comprehenders have typical word reading skill and intelligence but poorer than expected reading comprehension. While the prevalence of poor comprehenders is similar to that of poor decoders (individuals who have difficulty fluently converting written text into spoken language), less is known about the neurobiological substrates of poor comprehension. Extant studies have found small differences in grey matter volume between poor comprehenders and poor or typically reading peers. However, a detailed quantification of cortical morphometric features and white matter integrity remains unexplored.

Data from 2,100 children (1,200 with imaging data), aged 8-16 were analyzed to determine if there is a distinct neuroanatomy associated with poor reading comprehension. We computed grey matter volume, cortical thickness, and surface area, and white matter measures including mean diffusivity, fractional anisotropy, neurite orientation, and neurite density for poor comprehenders and compared to that of poor decoders and typical readers.

Results revealed small but widespread white matter differences, but no grey matter differences between poor comprehenders and other readers. Poor comprehenders showed decreased white matter integrity (increased mean diffusivity, decreased neurite density) in tracts previously associated with reading, including the Superior Longitudinal Fasciculus and Inferior Longitudinal Fasciculus, and in tracts that have been associated with cognitive performance such as the Uncinate Fasciculus.

These results suggest that diffuse structural connectivity differences may underlie reading comprehension weaknesses in the face of intact decoding skills. This is consistent with the behavioral profile of poor comprehenders who exhibit a broad pattern of subclinical impairments in language and integrative cognitive processes.

## Introduction

Reading is vital for both academic and personal success, yet there is substantial variation in reading comprehension with recent reports documenting poorer than expected reading comprehension performance among both children and adults in the US and beyond.^1–8^ Neurobiological studies of reading have been instrumental in documenting individual differences in single word reading and decoding, which has helped to isolate possible mechanisms of reading variation.^10–24^ However, there has been limited study of individual differences in reading comprehension. A small body of literature documents functional and structural differences between good and poor comprehenders; however, this literature is plagued by small sample sizes, variable classification of poor comprehenders, and inconsistent findings. The purpose of the current study is to provide the first large scale investigation of structural neuroanatomic characteristics of good and poor comprehenders as compared to poor decoding and typically reading peers.

It’s well understood that individuals can vary in their performance across multiple levels of reading skill, from single word decoding to text comprehension.^8,25–27^ While substantial research has focused on poor decoders/dyslexics and generally poor readers, relatively less work has focused on individuals with good single word reading skills but poor reading comprehension, so called “poor comprehenders” (PCs).^28–30^ In addition to poor reading comprehension relative to decoding, these readers tend to have sub-clinical weaknesses in oral language skills, including smaller vocabularies and weaker morphological and grammatical processing.^31–36^ PCs also display poorer than average performance on complex integrative tasks, such as inference making and comprehension monitoring^5,31,37–38^ and subtle processing weaknesses in domain-general skills such as executive functions and category learning.^9,30,39–41^ Across this literature, there is debate as to whether PCs are best characterized as having subclinical developmental language disorder or under a multifactorial model with both language and more domain-general cognitive difficulties that may stem from independent causal mechanisms.^30,39,42–46^ While neurobiological studies could help to shed light on this issue, at present there are only a few studies with inconsistent findings that have directly compared brain structure or function on PCs to that of typical readers and/or poor decoders.

Using matched group designs, two previous studies compared functional brain activation in PCs as compared to typical readers and/or poor decoders. A study by Cutting and colleagues^11^ found increased activation for PCs (n=12 adolescents) as compared to poor decoders (n=20 adolescents) during single word reading (lexical decision) in the left fusiform gyrus, left middle frontal and left cingulate gyrus. This is consistent with better single word reading performance in PCs relative to dyslexic readers.^11^ When compared to typical readers (n=19 adolescents), PCs had less activation in the right lingual gyrus and left cuneus for low frequency words only, and greater activation in the left inferior frontal gyrus (IFG) compared to right IFG. This is consistent with differential access to semantic information from visual information between PCs and typical readers. Connectivity analyses revealed that PCs had greater functional connectivity between left IFG and a number of cortical and subcortical regions including the left parahippocampal and hippocampal gyri and right thalamus, putamen, and middle frontal gyrus as compared to typical readers, suggesting less efficient or less automatic access of lexical semantic information in PCs relative to typical readers.^11^

A second study by Feng and colleagues^14^ used an auditory rhyme judgement task and found reduced activation for PCs (n=16 adolescents) relative to typical readers (n=102 adolescents) in the left middle frontal gyrus, IFG, insula, inferior temporal gyrus/middle occipital gyrus (fusiform gyrus), and anterior cingulate gyrus and increased activation in the right precentral, anterior cingulate, and supramarginal gyri, and left posterior cingulate, angular gyrus, middle frontal gyrus, and medial superior frontal gyrus.^14^ The differences observed here, with Chinese speakers, are in partial contrast to previous findings, which reveal typical rhyme processing for PCs.^47^ However, note that the PCs in this study had significantly poorer character naming than typical readers, and thus may not be comparable to most other studies of PCs, wherein word-level reading skills are matched.^14^ Moreover, psycholinguistic differences between the English and Chinese, including the use of tones to distinguish meaning in Chinese and the greater opacity of the Chinese writing system, complicate direct comparisons.

Other studies have investigated brain activation associated with reading comprehension skill during passage reading and/or listening using regression-based approaches which control for decoding skill and general cognitive skill. Aboud and colleagues^10^ used a passage reading task with 38 adolescents and found that those with higher reading comprehension scores showed increased activation in the left IFG extending into the left insula and dorsolateral prefrontal cortex. Additional positive correlations were found between reading comprehension and bilateral temporoparietal activation extending into the right ventral IFG, ventral insula, and left IFG as well as the default mode network.^10^ Using a similar approach, Ryherd and colleagues^13^ examined functional activation during passage reading and story listening in 32 adolescents. They found that those with better reading comprehension had increased activation across a broad network of regions important for language, including the left IFG, fusiform gyrus, middle temporal gyrus, and left cerebellum during both reading and listening. Whereas those with lower reading comprehension had greater activation in regions linked to attention and executive functions, including bilateral anterior and posterior cingulate, right middle frontal gyrus and right cerebellum.^13^

Although the functional tasks, languages/writing systems, and the methods used to identify the neural signatures of comprehension differed across these studies, better comprehenders tended to show greater activation during language and reading tasks, particularly in left hemisphere and bilateral regions that have been linked to language, attention, and memory, which may suggest greater inferencing, comprehension monitoring, and overall engagement with printed and spoken passages. However, some of these studies also observe brain regions with greater activation for PCs, typically associated with executive functions, though also in some regions linked to higher level language processing. Given that these studies focused on adolescents and adults, this overall pattern may represent a combination of atypical pathways for language and/or executive functions and compensation for weaknesses in the primary language processing systems. However, study differences in task, population, PC classification approach, and small samples preclude strong conclusions.

With respect to brain structure, studies have investigated grey matter volume in PC’s relative to typical readers and poor decoders. Using multivariate pattern analyses and a matched group design, Bailey and colleagues^9^ analyzed data from 41 adolescents and found reduced grey matter volume for PCs (n=11) relative to poor decoders (n=14) and typical readers (n=16) in a network of bilateral regions that subserve language and more general cognitive functions. Regions identified included right: superior/middle frontal gyrus, superior/inferior temporal gyrus, precentral gyrus, anterior cingulate, and the cuneus and cerebellum; and left: superior frontal gyrus, inferior/superior temporal gyrus, and middle occipital gyrus. No regions were found to be significantly larger for PCs relative to poor decoders and typical readers.^9^ Using a regression-based approach with 55 adolescents, Patael and colleagues^12^ attempted to identify regions associated with a discrepancy between reading comprehension and decoding. They found that grey matter volume in the left dorsolateral prefrontal cortex (dlPFC) was related to reading comprehension and the discrepancy between reading comprehension and decoding performance such that participants with higher reading comprehension performance relative to their decoding performance (unexpected good comprehenders) had increased grey matter volume in the left dlPFC as compared to those with relatively higher decoding as compared to their reading comprehension.^12^

While there are no studies of PCs that we know of that have examined white matter tractography, given inclusion of tractography data in the current study, we note two studies that have looked at correlations between reading comprehension and white matter integrity (without controlling for decoding or word level reading). Both studies found a positive correlation between reading comprehension and fractional anisotropy (FA) in the bilateral arcuate fasciculus (AF), inferior longitudinal fasciculus (ILF), and superior longitudinal fasciculus (SLF).^23–24^ While we use these studies to guide our predictions in the absence of white matter studies of PCs, it’s important to note that these same tracts have been similarly correlated with word level reading skills and even phonological processing and thus associations may not be specific to reading comprehension after controlling for decoding or word level reading skills.

Collectively, these structural findings are partially consistent with the functional findings and suggest broad associations between reading comprehension and brain anatomy in both language-associated and more domain-general processing linked regions. However, with only two relatively small studies of grey matter structure in PCs that used different analytic approaches and no white matter studies of PCs, this conclusion is tenuous and incomplete. White matter investigations may be particularly informative as reading comprehension is expected to require integration of information processed in multiple anatomically distinct regions, thus requiring cross regional communication. Hence, to improve understanding of the neurobiological basis of PCs and thus potentially shed light on the mechanisms of impairment, we used a large publicly available dataset to examine grey matter and white matter correlates of poor reading comprehension in the absence of poor decoding. We utilize three common classification approaches for identification of PCs and relevant controls to determine whether the classification approach would affect the regions or tracts identified. Based on aggregated findings from previous functional and structural neuroimaging studies of good and poor comprehenders, we made the following predictions: 1. PCs would show decreased grey matter morphometry in right- and left-hemisphere regions implicated in language and attention (e.g., right superior and middle frontal gyri, superior and inferior temporal gyri, precentral, postcentral, anterior cingulate, and lingual gyrus; left superior frontal gyrus, superior and inferior temporal gyri, middle occipital gyrus, inferior parietal lobule and dlPFC), and in the hippocampus as compared to all other readers.^9,14^ 2. PCs/unexpected PCs would have reduced white matter integrity (i.e., lower FA, higher MD) in the arcuate fasciculus (AF), superior longitudinal fasciculus (SLF), and inferior longitudinal fasciculus (ILF), bilaterally.^16,23–24^

## Materials and methods

### Participants

Behavioral data and neuroimaging scans were selected from 4,748 children, aged 5-21 years, from the Child Mind Institute Healthy Brain Network (CMI-HBN) biobank.^48^ All participants or their parents gave informed consent to participate in the study in accordance with the Declaration of Helsinki and the Chesapeake Institutional Review Board.^48^ To be included in the current study, participants had to be at least 8 years of age, have a nonverbal IQ score of ≥70, have data for all behavioral assessments of interest, and have no clinician-confirmed diagnosis of Autism Spectrum Disorder, Intellectual Disability, or Borderline Intellectual Functioning (full list of excluded disorders can be found in Supplementary Table 2). The final sample included 2,129 participants (age in years, *M* = 11.11, *SD=* 2.23), 1,270 of whom had 3T T1w MRI data available, with 903 participants also having diffusion data available.

### Brain Image Acquisition and Processing

All Healthy Brain Network Biobank brain imaging data used in this analysis (whole head) was acquired on 3 Tesla scanners using protocols detailed elsewhere.^48,49^ T1w images were acquired with a TR=2500ms, echo time =3.15ms, voxel size=0.8 mm (isometric), and flip angle = 8°. Diffusion weighted images (DWI) were obtained in 64 directions with bvals =0, 1000, 2000sec/mm^2^ and isotropic voxels of 1.8mm. Raw images were downloaded in BIDS format and were processed in-house.

### MRI Imaging Quality Control and Preprocessing of Images

T1w image quality was assessed using MRIQC (version 22.0.6) and then processed using the FreeSurfer (version 7.1.1) automated pipeline for surface-based cortical reconstruction and volumetric segmentation.^50–51^ The Freesurfer image analysis suite is documented and freely available for download online. The technical details of these procedures are described in prior work.^51–64^ Briefly, this processing includes motion correction and averaging of volumetric T1 weighted images,^63^ removal of non-brain tissue using a hybrid watershed/surface deformation procedure,^64^ automated Talairach transformation, segmentation of the subcortical white matter and deep grey matter volumetric structures^55,58^ intensity normalization,^65^ tessellation of the grey matter white matter boundary, automated topology correction,^54,66^ and surface deformation following intensity gradients to optimally place the grey/white and grey/cerebrospinal fluid borders at the location where the greatest shift in intensity defines the transition to the other tissue class.^51–53^ Once the cortical models are complete, a number of deformable procedures were performed for further data processing and analysis including surface inflation,^53^ registration to a spherical atlas which is based on individual cortical folding patterns to match cortical geometry across subjects,^56^ parcellation of the cerebral cortex into units with respect to gyral and sulcal structure,^59,67^ and creation of a variety of surface based data including maps of curvature and sulcal depth. Processed data was mapped to the Destrieux atlas to extract cortical thickness, surface area and volume, and a whole brain approach was used for all subsequent analyses, testing each of the 74 regions in each hemisphere for 1,270 participants.^51^

Diffusion MRI (dMRI) images were processed using Functional Magnetic Resonance Imaging of the Brain Software Library (FMRIB FSL, version 6.1.0).^68–69^ The Quality Assessment of all dMRI data (QUAD) toolbox in FSL was used to estimate multiple quality metrics including contrast to noise ratio (CNR) for all DTI data.^69^ Standard pipelines and general best practices for preprocessing diffusion data in FSL are well outlined elsewhere^68,70^ and the specifics of diffusion processing for the data in this project are similarly outlined elsewhere.^17,71–72^ In short, data was preprocessed for artifact correction including susceptibility, eddy current correction, and correcting for head movement. The Brain Extraction Toolkit (BET) was then used to create individual masks for each brain, isolating the brain from the skull. FSL’s diffusion tensor modeling tool, DTIfit, was used to obtain diffusion measures including fractional anisotropy (FA) and mean diffusivity (MD). The crossing fibers distribution was estimated using BEDPOSTX and the probability of major fiber directions were calculated. A multi-fiber model was fit at each voxel for tracing fibers through regions of crossing or complexity. XTRACT was used to compute probabilistic tractography (curvature threshold of ±80°, max streamline steps: 2000) in each participant’s native space to obtain 23 major fiber tracts within the human brain. The normalized fiber probability distribution obtained was then thresholded and binarized using fslmaths to get a tract mask. The mask generated was inspected for accuracy and the value was adapted accordingly to ensure correct segmentation. The obtained binary mask was then multiplied by the subject’s NODDI and diffusion maps to get a tract specific distribution of those values. All tracts, along with their abbreviations, are listed in Supplementary table 1.

Additionally, to better characterize the white matter microstructure, the neurite orientation dispersion and density imaging (NODDI) model was fit using the NODDI toolbox.^73^ NODDI is a multi-compartment, non-Gaussian, biophysical tissue model that can quantitatively evaluate specific microstructural changes to distinguish between three microstructural environments: intracellular, extracellular and CSF compartments.^73^ The amount of diffusion in each of these individual compartments allows for the estimation of density and orientation distribution of axons and dendrites, collectively termed neurites. NODDI coefficients neurite orientation and dispersion index (ODI) and neurite density index (NDI) were mapped along 23 fiber tracts and mean values for each tract were used for all subsequent analyses.

### Behavioral Measures

To categorize children into poor comprehender, poor decoder, or typical reader groups, behavioral assessments of reading and language were used. Reading comprehension was measured through the Wechsler Individual Achievement Test-3^rd^ edition (WIAT) reading comprehension subtest during which participants read short expository and narrative passages and answer comprehension probe questions.^74^ Decoding was measured using the Test of Word Reading Efficiency-2^nd^ edition (TOWRE) pseudoword decoding efficiency subtest during which participants read as many pronounceable English pseudowords as possible in 45 seconds.^75^ Receptive vocabulary, commonly used as a proxy measure of listening comprehension in the PC literature, was measured using the WIAT-3^rd^ edition receptive vocabulary subtest, during which participants indicate which of four pictures best illustrates a word they hear.^74^ Lastly, the Wechsler Intelligence Scales for Children- 5^th^ edition (WISC) was used to measure nonverbal IQ which included block design, matrix reasoning, coding, figure weights, visual puzzles, and picture span subtests which were summed and standardized to create the nonverbal index score.^76^ Raw behavioral assessment scores, along with demographic information including age and sex, were then used to determine group classification and were used as covariates in analyses (see Table 1).

**Table 1:** Demographic information for all participants and classification methods

|  | N (Female) | Age<sup>a</sup> | Reading Comprehension<sup>b</sup> | Decoding<sup>b</sup> | Oral Vocabulary<sup>b</sup> | Nonverbal IQ |
| --- | --- | --- | --- | --- | --- | --- |
| **Grey Matter Analyses** |  |  |  |  |  |  |
| **Cutoff Classification** |  |  |  |  |  |  |
| Poor Comprehenders | 58 (13) | 11.57 (2.04) | 85.50 (6.00) | 106.43 (6.23) | 95.83 (17.81) | 90.00 (11.76) |
| Poor Decoders | 58 (16) | 10.82 (1.78) | 108.64 (10.24) | 82.41 (6.24) | 113.22 (9.12) | 97.93 (12.12) |
| Typical Readers | 58 (13) | 11.38 (2.05) | 109.45 (10.48) | 110.22 (8.23) | 114.48 (9.38) | 92.66 (10.42) |
| **Mixed Regression** |  |  |  |  |  |  |
| Unexpected Poor Comprehenders | 254 (81) | 11.00 (2.05) | 86.37 (9.13) | 94.11 (16.82) | 104.69 (14.77) | 97.28 (14.62) |
| Expected Comprehenders | 254 (99) | 11.15 (2.24) | 101.67 (9.18) | 99.41 (16.46) | 108.50 (14.38) | 102.81 (15.85) |
| Unexpected Good Comprehenders | 254 (103) | 10.95 (2.28) | 112.91 (15.41) | 92.90 (14.44) | 103.53 (13.51) | 97.16 (14.16) |
| **Regression** |  |  |  |  |  |  |
| Full Sample | 1270 (480) | 10.99 (2.16) | 100.73 (14.76) | 96.86 (16.34) | 105.66 (15.02) | 100.01 (15.12) |
| **White Matter Analyses** |  |  |  |  |  |  |
| **Cutoff Classification** |  |  |  |  |  |  |
| Poor Comprehenders | 26 (7) | 11.60 (2.12) | 85.27 (4.80) | 106.88 (5.39) | 96.62 (18.51) | 88.27 (11.29) |
| Poor Decoders | 26 (7) | 11.12 (1.73) | 106.81 (8.47) | 80.62 (7.58) | 110.42 (7.30) | 92.35 (11.87) |
| Typical Readers | 26 (7) | 11.53 (2.21) | 111.42 (12.56) | 111.46 (8.59) | 112.50 (9.48) | 89.15 (10.62) |
| **Mixed Regression** |  |  |  |  |  |  |
| Unexpected Poor Comprehenders | 181 (63) | 10.77 (1.97) | 86.39 (9.85) | 92.41 (17.98) | 103.81 (14.51) | 96.42 (15.61) |
| Expected Comprehenders | 181 (69) | 11.02 (2.21) | 103.31 (10.60) | 99.30 (16.94) | 107.50 (14.63) | 103.12 (15.57) |
| Unexpected Good Comprehenders | 181 (65) | 11.02 (2.29) | 113.52 (15.54) | 92.65 (13.94) | 103.49 (13.79) | 95.89 (13.96) |
| **Regression** |  |  |  |  |  |  |
| Full Sample | 903 (336) | 11.05 (2.18) | 102.11 (15.58) | 96.83 (16.44) | 105.57 (14.85) | 100.08 (15.35) |

<sup>a</sup>In all relevant cases mean is reported with standard deviation in parenthesis.
<sup>b</sup>All behavioral scores reported are standardized scores for interpretability, while raw scores were used in all models. All behavioral tests have a test mean of 100 and standard deviation of 15.

### Group Classification

Three widely used classification approaches were utilized for assessing poor comprehenders: a) a cutoff classification model which identifies PCs based on standard scores indicating poor reading comprehension and good decoding and compares them to age, sex, and nonverbal IQ matched poor decoders and good readers; b) a mixed classification model which uses regression to predict reading comprehension from decoding, vocabulary, age, sex, and nonverbal IQ, and divides the residuals into unexpected poor comprehender, unexpected good comprehender, and expected comprehender groups; and, c) a continuous approach which jettisons groups altogether and uses the regression residuals from b) to predict the brain measures (See Figure 1).

**Figure 1:**
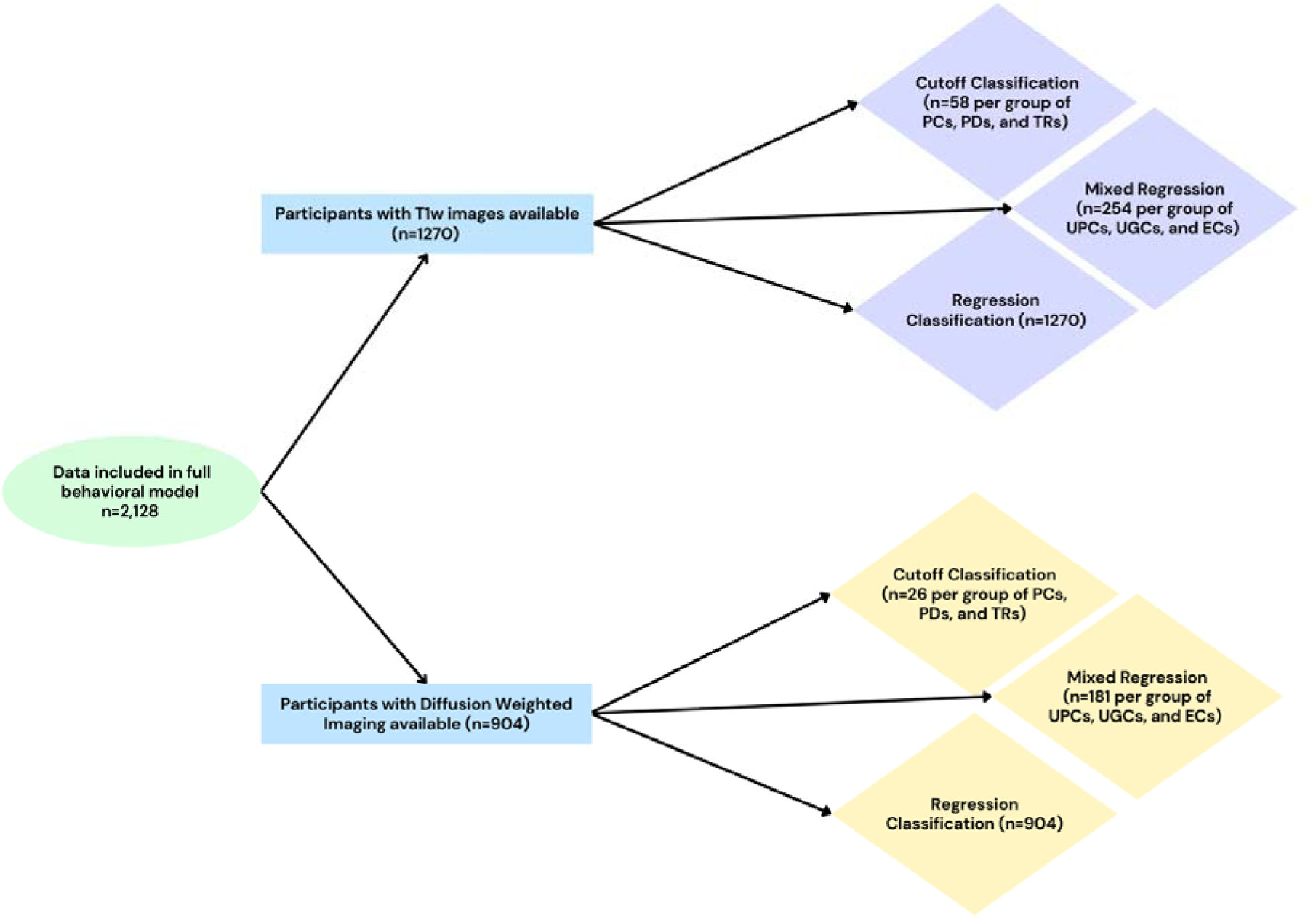
Graphical representation of data selection processes. Figure depicts the data selection and group classification processes used in the current analyses. PCs- poor comprehenders; PDs- poor decoders; TRs- typical readers; UPCs- unexpected poor comprehenders, UGCs- unexpected good comprehenders, ECs- expected comprehenders

### Cutoff Classification Model

Consistent with previous literature, this model identifies PCs (*n*=58 for grey matter analysis, *n*=26 for white matter analysis) as those with typical decoding (decoding standard score of ≥100), and lower than average reading comprehension (reading comprehension standard score of ≤90). We then identified age, sex, and nonverbal IQ matched groups of poor decoders (*n*=58 grey matter, 26 white matter, who had decoding standard scores of ≤90 and reading and oral language standard scores ≥100) and typical readers (*n*=58 grey matter, 26 white matter who had decoding, reading, and oral language standard scores of ≥100) using the *MatchIt!* R package (version 4.5.5, optimal method, Mahlanobis distance).^77^ Grey and white matter groups were matched independently. One outlier was identified in the PC grey matter group, with a reading comprehension score which was 10 points lower than the next poorest comprehending peer, though no other behavioral or brain measures were found to be significantly different from the group. All models were run with and without the outlier to assess possible skew in results.

### Mixed Classification Model

This model uses regression to predict reading comprehension from decoding, oral language abilities, and control variables (age, sex, and non-verbal IQ), operationalizing the Simple View of Reading.^8^ All participants with behavioral data (*n*=2,129) were included in this model to provide the most representative distribution of reading scores available given our inclusion criteria. After modeling the relationship between reading comprehension and other variables, residual scores were used to define group membership and the lowest, middle, and highest 20% of residual scores with imaging data available were retained for analysis.^78^ Participants with residual scores in the 40^th^ to 60^th^ percentiles (which were generally close to zero, indicating little difference between predicted and expected reading comprehension) formed the “expected comprehender group” (EC, *n*=254 grey matter, 181 white matter). Those with positive residual scores in the 80^th^ to 100^th^ percentiles (indicating better than predicted reading comprehension) formed the “unexpected good comprehender group” (UGC, *n*=254 grey matter, 181 white matter). Participants with negative residual scores in the 0^th^ to 20^th^ percentiles (indicating worse than predicted reading comprehension) formed the “unexpected poor comprehender group” (UPC, *n*=254 grey matter, 181 white matter). No outliers were identified in these groups.

### Regression Classification Model

In this final model residual scores from the mixed classification model detailed above were directly used to predict brain measures (*n*=1,270 grey matter, 903 white matter). See Table 1 for a description of participants across all classification models.

### Statistical Analysis

Across both grey and white matter, linear models were estimated for each classification approach and for each brain metric across the whole brain. Age, nonverbal IQ scores, and imaging quality measures (CNR for both grey and white matter) were centered in all analyses. All models used group status (PC, typical readers, etc.) or residual score and age, sex, nonverbal IQ, and MRI quality control measures as covariates to estimate brain structure. In models using group status, group was dummy coded with PCs being coded as 1 in all cases. All models were fit in R (version 4.3.2) using the *lm* function and corrected for multiple comparisons using the Benjamini-Hochberg false discovery rate correction.^79–80^ Corrections were applied to each dependent variable of interest independently for each classification method across both grey and white matter analyses for each contrast (PCs compared to poor decoders, typical readers, and all other readers combined, etc.).

## Results

### White Matter

#### Cutoff Classification Method

No significant differences in white matter structure (FA, MD, NDI, ODI) were identified by the cutoff classification models.

#### Mixed Regression Classification Method

In the mixed regression classification method, small but systematic differences in white matter structure were found between unexpected poor comprehenders and other readers. Specifically, unexpected poor comprehenders showed increased MD as compared to both expected comprehenders and unexpected good comprehenders across a number of left and right hemisphere tracts (all results *p*<.05 after FDR correction, see Table 2 for a full list of tracts which differed between groups). These differences were found in tracts that have commonly been linked to variation in reading and language, including the bilateral SLF and ILF, bilateral arcuate fasciculus, bilateral inferior fronto-occipital fasciculus, and middle longitudinal fasciculus. Increased MD for unexpected poor comprehenders (UPCs) as compared to expected comprehenders and unexpected good comprehenders was also found in tracts that have been correlated with primarily domain-general processes (i.e., attention), including the anterior commissure, bilateral fornix, bilateral uncinate fasciculus, and others.

**Table 2:** Significant differences in white matter structure by contrast

| Mean Diffusivity | Significant Contrasts<sup>a</sup> |
| --- | --- |
| Anterior Commissure | UPC vs EC, UPC vs UGC, UPC vs All |
| L Arcuate Fasciculus | UPC vs EC, UPC vs UGC, UPC vs All |
| R Arcuate Fasciculus | UPC vs EC, UPC vs UGC, UPC vs All |
| L Acoustic Radiation | UPC vs UGC, UPC vs All |
| L Anterior Thalamic Radiation | UPC vs EC, UPC vs UGC, UPC vs All |
| R Anterior Thalamic Radiation | UPC vs EC, UPC vs UGC, UPC vs All |
| L Dorsal Cingulum | UPC vs EC, UPC vs UGC, UPC vs All |
| R Dorsal Cingulum | UPC vs EC, UPC vs UGC, UPC vs All, Residuals |
| L Perigenual Cingulum | UPC vs EC, UPC vs UGC, UPC vs All, Residuals |
| R Perigenual Cingulum | UPC vs EC, UPC vs UGC, UPC vs All, Residuals |
| L Temporal Cingulum | UPC vs EC, UPC vs UGC, UPC vs All |
| R Temporal Cingulum | UPC vs EC, UPC vs UGC, UPC vs All |
| L Frontal Aslant | UPC vs EC, UPC vs UGC, UPC vs All |
| R Frontal Aslant | UPC vs EC, UPC vs UGC, UPC vs All |
| Forceps Major | UPC vs UGC, UPC vs All |
| Forceps Minor | UPC vs EC, UPC vs UGC, UPC vs All |
| L Fornix | UPC vs EC, UPC vs UGC, UPC vs All |
| R Fornix | UPC vs EC, UPC vs UGC, UPC vs All, Residuals |
| L Inferior Fronto-Occipital Fasciculus | UPC vs EC, UPC vs UGC, UPC vs All |
| R Inferior Fronto-Occipital Fasciculus | UPC vs EC, UPC vs UGC, UPC vs All |
| L Inferior Longitudinal Fasciculus | UPC vs UGC |
| R Inferior Longitudinal Fasciculus | UPC vs EC, UPC vs UGC, UPC vs All |
| Middle Cerebral Peduncle | UPC vs EC, UPC vs All |
| L Middle Longitudinal Fasciculus | UPC vs EC, UPC vs UGC, UPC vs All |
| R Middle Longitudinal Fasciculus | UPC vs EC, UPC vs UGC, UPC vs All |
| L Optic Radiation | UPC vs EC, UPC vs UGC, UPC vs All |
| R Optic Radiation | UPC vs EC, UPC vs UGC, UPC vs All |
| L Superior Longitudinal Fasciculus (average) | UPC vs EC, UPC vs UGC, UPC vs All |
| R Superior Longitudinal Fasciculus (average) | UPC vs EC, UPC vs UGC, UPC vs All |
| L Superior Thalamic Radiation | UPC vs EC, UPC vs UGC, UPC vs All |
| R Superior Thalamic Radiation | UPC vs UGC |
| L Uncinate Fasciculus | UPC vs EC, UPC vs UGC, UPC vs All |
| R Uncinate Fasciculus | UPC vs EC, UPC vs UGC, UPC vs All |
| L Vertical Occipital Fasciculus | UPC vs UGC, UPC vs All |
| R Vertical Occipital Fasciculus | UPC vs UGC, UPC vs All |
| **Neurite Density Index** |  |
| L Anterior Thalamic Radiation | UPC vs EC, UPC vs All |
| L Dorsal Cingulum | UPC vs EC, UPC vs UGC, UPC vs All |
| R Dorsal Cingulum | UPC vs EC, UPC vs UGC, UPC vs All, Residuals |
| L Perigenual Cingulum | UPC vs EC, UPC vs UGC, UPC vs All, Residuals |
| R Perigenual Cingulum | UPC vs EC, UPC vs UGC, UPC vs All, Residuals |
| R Temporal Cingulum | UPC vs UGC |
| L Frontal Aslant | UPC vs UGC, UPC vs All |
| R Frontal Aslant | UPC vs EC, UPC vs All |
| R Fornix | UPC vs All |
| R Inferior Fronto-Occipital Fasciculus | UPC vs EC, UPC vs UGC, UPC vs All |
| R Inferior Longitudinal Fasciculus | UPC vs UGC |
| L Middle Longitudinal Fasciculus | UPC vs UGC |
| L Superior Longitudinal Fasciculus (average) | UPC vs EC, UPC vs All |
| L Uncinate Fasciculus | UPC vs EC, UPC vs UGC, UPC vs All |

All white matter results which remained significant ( $p < 0.05$ ) after Benjamini-Hochberg false discovery rate multiple comparisons correction.
<sup>a</sup>UPC= Unexpected Poor Comprehender, EC= Expected Comprehender, UGC= Unexpected Good Comprehender, All = Combined Expected Comprehender and Unexpected Good Comprehender group, Residuals= Differences seen in residuals used continuously in the regression classification.

Similarly, UPCs were found to have decreased neurite density index (NDI) as compared to both expected comprehenders and unexpected good comprehenders in a more limited number of tracts (see Table 2). Like with mean diffusivity, some of these differences were found in tracts that connect regions involved with language processing, including the left SLF and right ILF. Additionally, a small number of tracts that have been linked to domain-general processes were implicated including the left anterior thalamic radiation and uncinate, right fornix and temporal cingulum, and bilateral dorsal and perigenual cingulum and frontal aslant.

No statistically significant differences were observed between unexpected poor comprehenders and peers in FA or ODI.

#### Regression Classification Method

In the regression classification, participants with more negative residual scores (those whose comprehension was poorer than expected) were found to have increased MD as compared to those with more positive residual scores in the right dorsal cingulum, fornix, and bilateral perigenual cingulum.

Two of these results were mirrored in the NODDI measures, which indicated that those with more negative residual scores had decreased NDI as compared to those with more positive residual scores in the right dorsal cingulate and bilateral perigenual cingulum.

### Grey Matter Analyses

No statistically significant results were observed after correction for multiple comparisons for any classification models. However, we did find a number of nominally significant results which are consistent with prior literature and thus discussed briefly below (and see Supplementary Table 3 for full details of these results).

## Discussion

The current study sought to determine whether PCs differ from typical readers or poor decoders in their brain structure using a large, diverse sample. This study is the first that we know of to examine and compare white matter microstructural integrity of these groups, and the first to compare these groups on cortical morphometric measures like surface area and cortical thickness. Furthermore, this study also compares whether classification approach affects the regions or tracts associated with poor comprehenders or that differ between groups.

Our results reveal widespread but small differences in white matter tract integrity between PCs, poor decoders, and typical readers. Specifically, effects that differentiated the groups were found bilaterally, in tracts that have been linked to variation in language ability as well as in tracts that have been linked to domain-general processes (Figure 2). Some of these findings (discussed in detail below) are consistent with existing MRI studies of reading comprehension, and thus our hypotheses, while others were observed for the first time. Additionally, our study provides evidence that diffusivity (MD) and neurite density (NDI) may be more sensitive to comprehension related variation in white matter structure than anisotropy, possibly relating it to the impact of neurodevelopmental processes like neuronal migration, axon guidance, myelination, and synaptic pruning.

**Figure 2:**
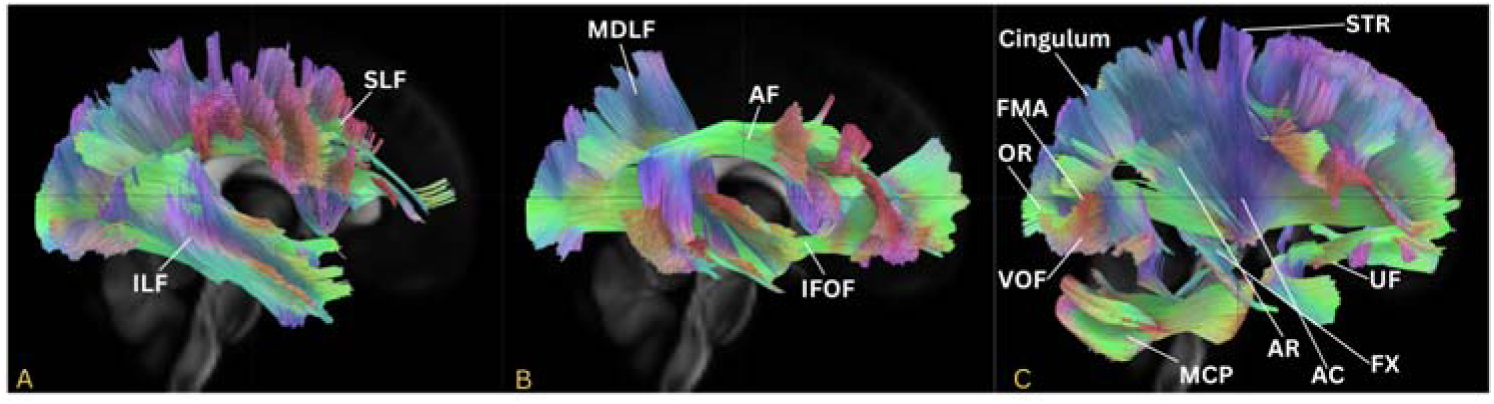
White matter tracts with significant differences in UPCs versus other readers Panel A shows hypothesized white matter tracts with significantly worse integrity in UPCs than other readers. Panel B shows language-related tracts with significantly worse integrity in UPCs than other readers. Panel C shows domain-general tracts with significantly worse integrity in UPCs than other readers. All tract abbreviations can be found in Supplementary Table 1.

### White Matter Analyses

While the literature on white matter in poor comprehenders is negligible, some of our findings are consistent with findings from studies which have examined variation in reading comprehension in typical readers.^23–24^ Specifically, we found reduced integrity for unexpected PCs (increased MD and decreased NDI) as compared to expected comprehenders and unexpected good comprehenders in the bilateral SLF and ILF, as well as in the bilateral arcuate fasciculus. While previous studies only considered fractional anisotropy, a metric for which we found no significant differences, our results are regionally and directionally consistent. We also observed reduced integrity for PCs (increased MD and decreased NDI) as compared to expected comprehenders and unexpected good comprehenders in two tracts that have previously been associated with language or reading, but not with reading comprehension specifically: the inferior fronto-occipital fasciculus, and the middle longitudinal fasciculus. In previous studies, poorer integrity in these tracts was correlated with worse word level reading skills or language skills.^17,81–85.^ Specifically, the inferior fronto-occipital fasciculus, especially in the left hemisphere, is thought to act as a ventral pathway supporting orthographic awareness and word reading, with tract integrity predicting more accurate performance on an orthographic awareness task in adult readers with varying reading abilities^82–83^ and the middle longitudinal fasciculus has recently been linked to impaired semantic processing in aphasics, and has been correlated with performance on a variety of expressive and receptive language tasks after brain injury (i.e., the Progressive Aphasia Severity Scale).^81,84^

In addition to tracts commonly associated with language and reading, we also found reduced integrity (increased MD and decreased NDI) in unexpected PCs in several tracts that have been implicated in both language *and* more general cognitive function including the bilateral uncinate fasciculus, frontal aslant tract, and acoustic radiation. A recent review article posits that the uncinate fasciculus may play a role in semantic processing, emotional processing, and in learning, including correlations with primary progressive aphasia and language performance in the disorder.^85^ Indeed, there is mounting evidence that uncinate fasciculus integrity is correlated with semantic tasks.^85^ Similarly, the frontal aslant tract is hypothesized to play a role both in language and in general cognition and is hypothesized to support language learning, language comprehension, and fluency and/or stuttering.^85^ A recent review article has highlighted that left frontal aslant tract integrity is typically correlated with stutter severity as well as more general tasks like executive functions, particularly inhibitory control.^85–86^ Finally, the left acoustic radiation connects thalamic and temporal areas and is thought to relay acoustic information, but limited research suggests that it may also be important for more general language and auditory processing.^87^

We also observed reduced integrity for unexpected PCs relative to expected comprehenders, unexpected good comprehenders, and a combined group of expected and unexpected good comprehenders in tracts linked to visual processing, including the bilateral optic radiation, vertical occipital fasciculus, and superior thalamic radiation. The optic radiations are primarily linked to vision and visual tasks.^88–89^ The vertical occipital fasciculus has been hypothesized to connect the dual visual stream and also play a role in integrating visual information, but tractography studies correlated to these skills are still developing.^90^ The superior thalamic radiation has not been characterized as extensively as many other tracts, but one recent study found that FA of the tract was related to eye movement characteristics in participants with attenuated psychosis syndrome.^91^ This suggests that PCs may have trouble efficiently integrating visual information, which could cause downstream effects in language and/or reading.

The final set of tracts identified here as having reduced integrity in unexpected PCs relative to expected comprehenders, unexpected good comprehenders, and a combined group of expected and unexpected good comprehenders have been associated with memory and information integration processes. These tracts include the fornix, anterior commissure, cingulum, and forceps major and minor. The fornix has been is predictive of cognitive impairment as well as being related to memory in young children.^92–94^ The anterior commissure has been correlated with primarily integrative tasks, such as the flow of information across the hemispheres.^95^ Like the anterior commissure, the cingulum is thought to primarily support integrative tasks, as well as executive functions including some evidence that those with reduced integrity in the cingulum perform worse on executive function tasks than those with greater integrity.^96–97^ While somewhat difficult to characterize the specific functions, the forceps major has been related to delayed memory in those with mood disorders while the forceps minor has been related to both attention deficits and bilingual status (possibly due to differences in task switching).^98–102^ These results provide further evidence that efficient information integration in many domains may be a weakness for poor comprehenders.

In addition to tracts that met the more stringent FDR corrected significance threshold, we found a small number of nominally significant differences in ODI that were hypothesized or mirrored the tracts discussed above. Nominally significant findings were found in the unexpected poor comprehenders versus all other readers (both expected comprehenders and unexpected good comprehenders) contrast of the mixed regression analysis, and reflected decreased ODI for unexpected PCs as compared to all other readers in the left anterior thalamic radiation, dorsal cingulum, superior longitudinal fasciculus, and bilateral fornix. Tthese results are directionally consistent with MD and NDI findings, potentially adding evidence for atypical tractography in PCs across these tracts.

Together, these tracts are thought to support numerous functions that relate to reading comprehension, including language, visual and auditory processing, memory, executive function, and interplay between the hemispheres. Generally, these results indicate that differences in white matter tract microstructure in poor comprehenders in our sample are found in a broad, diffuse, and bilateral set of tracts, supporting the hypothesis that PCs may have broad yet subtle differences in reading-related subskills which may be best characterized neuroanatomically through differences in connective (white matter) tissue.

### Grey Matter Analyses

In this study, we found no statistically significant differences in grey matter structure at our multiple comparisons corrected threshold. However, we found a number of nominally significant findings that replicate previous associations in the literature. These nominally significant findings were statistically significant before multiple comparisons correction but were non-significant (all p-values>0.10 after correction) after FDR correction. These findings include reduced grey matter volume in the right cerebellum for poor comprehenders vs typically developing readers in the cutoff classification and compared to unexpected good comprehenders in the mixed regression classification. Previous work has also found reduced right cerebellum volume in poor comprehenders as compared to both poor decoders and typical readers.^9^ Additionally, mirroring previous structural findings from the PC literature, unexpected PCs were found to have reduced cortical surface area in the bilateral precentral gyrus and right anterior cingulate as compared to unexpected good comprehenders in the mixed regression classification. Cortical surface area differences in the right precentral gyrus were also nominally significant in the regression classification model, in which those with more negative residuals had less cortical extent in the right precentral gyrus than those with more positive residuals. Lastly, mirroring functional differences seen in the previous literature, PCs were found to have less grey matter volume in the right amygdala as compared to all other readers in the cutoff classification method (with and without the outlier). Similarly, unexpected PCs were found to have reduced cortical surface area in the left cingulate and angular gyri and right anterior cingulate, supramarginal, and precentral gyri as compared to other readers and increased cortical thickness in the left middle frontal gyrus as compared to other readers, all areas which have been implicated in functional studies of PCs as compared to other readers. Other nominally significant findings are outlined in the supplementary materials (Supplementary Table 3).

With respect to differences between our study findings and findings from the two other grey matter studies of PCs in which more regions passed the threshold for statistical significance, we note two possible explanations. One possible explanation is differences in sample characteristics and size – our sample is larger and more diverse than most. Indeed, previous research suggests that reading related covariance in brain structure is more likely to be observed in homogeneous samples when variance in other environment-level and person-level factors is low^103^ and that replication attempts with brain wide association studies (BWAS) with larger samples are likely to yield fewer and smaller effects as they get closer to approximating the true effect sizes in the broader population, which are likely to be small for complex traits like reading.^104^ Second, it’s worth noting that our whole brain approach with ∼148 regions per brain metric required a conservative correction for multiple comparisons – if we had used an ROI based approach utilizing only those regions identified in one of the previous two grey matter studies of PCs, some of these regions (noted above and below) would have reached statistical significance (Supplementary Materials Table 3 notes regions which replicate previous findings). Thus, although nominally significant results should be interpreted with caution, together with previous findings they lend support for reading comprehension specific associations in the right cerebellum, amygdala, anterior cingulate, and supramarginal gyri, left cingulate gyrus, and bilateral precentral gyri.

### Classification Comparison

One sub-goal of the current study was to compare results across classification models. While expected patterns in behavior were generally observed across all approaches with PCs and UPCs with more similar IQ and decoding, there were a few differences in how distinct subtypes of readers were from each other across the approaches. Notably, the cutoff approach maximizes the difference between reading comprehension and decoding for the PCs and PDs (in opposite directions) while matching on other elements to each other and typical readers, whereas the mixed regression classification tends to produce less distinct UPCs and UGCs in terms of reading comprehension and decoding (see Supplementary Table 6). While these differences are small, they could contribute to the differences in the PC associated brain profiles observed across these two approaches.

In the white matter analyses, we observed many more effects which passed the FDR threshold for statistical significance in the mixed classification models relative to the regression classification models, and no tracts reached statistical significance in the cutoff classification models. Notably, all tracts which were statistically significant in the regression models were also significant in the mixed classification models. This pattern isn’t surprising given how these different approaches handle cases. The cutoff classification approach creates distinct groups, but yields small sample sizes per group giving it limited statistical power relative to the other two approaches. The mixed classification approach creates distinct groups and yields large sample sizes giving it good statistical power. The regression approach is a continuous approach which uses the same residuals as the mixed classification approach, but doesn’t eliminate the 20% of residuals that fall in between the expected comprehender and two unexpected comprehender groups, thus it has the largest sample size.

However, including all cases and assuming a linear relationship may have reduced power if the nature of the relationship between brain and behavior varies across the distribution. If participants in the outer quintiles – unexpected poor comprehenders and unexpected good comprehenders – have less extreme morphometric measures than would be predicted from their residuals, a model which assumes that a given difference in residuals predicts a specific (linear) change in these measures would not capture that fact, making it less likely to yield a statistically significant result (see Supplementary Figure 1 for exemplary graphs). Consistent with this largely power based explanation, examination of the coefficients by tract and metric indicates similarity in relative effect strength across all three approaches, with greater similarity between the two residual based approaches.

## Conclusion

In conclusion, our results suggest that in terms of brain structure, reading comprehension specific difficulties are undergirded by a diffuse network of regions and tracts that contribute to language processing, memory, emotion, and executive functions. These neurobiological findings are consistent with the broad but subtle behavioral impairments seen in PCs, including those consistently observed for complex integrative behaviors like comprehension monitoring and inference making. Our findings provide partial replication of previous grey matter studies of PCs with a large and diverse sample and further extend upon those studies by providing complementary white matter findings. Our findings also support a multifactorial model of poor comprehenders that includes difficulties in language and more domain-general processes. Future research should expand upon this work by examining functional correlates in a large and diverse sample.

## Data availability

All data and code used for this project, along with project preregistration, are publicly available in an OSF repository: Link to be inserted after review

## Supplementary material

Supplementary material is available online.

## Supplementary Material

### White matter tracts and abbreviations

**Supplemental Table 1:** White matter tracts and abbreviations

| Tract | Abbreviation |
| --- | --- |
| Anterior Commissure | AC |
| Arcuate Fasciculus (bilateral) | Left AF, Right AF |
| Acoustic Radiation (bilateral) | Left AR, Right AR |
| Anterior Thalamic Radiation (bilateral) | Left ATR, Right ATR |
| Dorsal Cingulum (bilateral) | Left CBD, Right CBD |
| Peri-genual Cingulum (bilateral) | Left CBP, Right CBP |
| Temporal Cingulum (bilateral) | Left CBT, Right CBT |
| Corticospinal Tract (bilateral) | Left CST, Right CST |
| Frontal Aslant Tract | Left FAT, Right FAT |
| Forceps Major | FMA |
| Forceps Minor | FMI |
| Fornix (bilateral) | Left FX, Right FX |
| Inferior Longitudinal Fasciculus (bilateral) | Left ILF, Right ILF |
| Inferior Fronto-Occipital Fasciculus (bilateral) | Left IFOF, Right IFOF |
| Middle Cerebellar Peduncle | MCP |
| Middle Longitudinal Fasciculus (bilateral) | Left MDLF, Right MDLF |
| Optic Radiation (bilateral) | Left OR, Right OR |
| Superior Longitudinal Fasciculus (bilateral)<sup>a</sup> | Left SLF, Right SLF |
| Superior Thalamic Radiation (bilateral) | Left STR, Right STR |
| Uncinate Fasciculus (bilateral) | Left UF, Right UF |
| Vertical Occipital Fasciculus (bilateral) | Left VOF, Right VOF |

<sup>a</sup>In this study, the three segments of the SLF were averaged in each hemisphere to create a single SLF variable per measure used for all analyses.

### Participants excluded for disorders

In this study, participants with a predefined set of diagnoses and disorders were excluded from all analyses. In total, 228 children who met other inclusion criteria were excluded because of a diagnosis.

**Supplemental Table 2:** Participants removed from analyses due to clinician confirmed diagnoses.

| **Clinician Confirmed or Suspected Diagnosis** | **Number of Participants Removed** |
| --- | --- |
| Autism Spectrum Disorder | 170 |
| Borderline Intellectual Functioning | 19 |
| Speech Sound Disorder | 15 |
| Pragmatic Communication Disorder | 10 |
| Intellectual Disability- Mild | 10 |
| Selective Mutism | 6 |
| Fetal Alcohol Related Neurodevelopmental Delay | 1 |
| *Total Participants Removed for Diagnosis* | 228 |

### Comparisons run for all imaging models

In the current study, the following comparisons were conducted for both grey and white matter analyses: in the cutoff classification model, poor comprehenders (binary coded as 1) were compared to poor decoders, typical readers, and poor decoders and typical readers combined (all binary coded as 0); in the mixed regression classification, unexpected poor comprehenders (binary coded as 1) were compared to unexpected good comprehenders, expected comprehenders, and unexpected good comprehenders and expected comprehenders combined (all binary coded as 0); in the regression classification, residuals were used to predict changes in brain structure continuously with unexpected poor comprehenders having lower residual values than expected comprehending or unexpected good comprehending peers.

### Supplementary grey matter results

In this study, there were no grey matter results which survived corrections due to multiple comparisons. This is likely due to the large number of comparisons (74 per hemisphere). There were, however, a number of nominally significant (significant before multiple comparisons corrections) findings of note. Poor comprehenders were coded as 1 in all group analyses and had more negative residual scores in regression classification models. Hypothesized findings are italicized.

**Supplemental Table 3:**
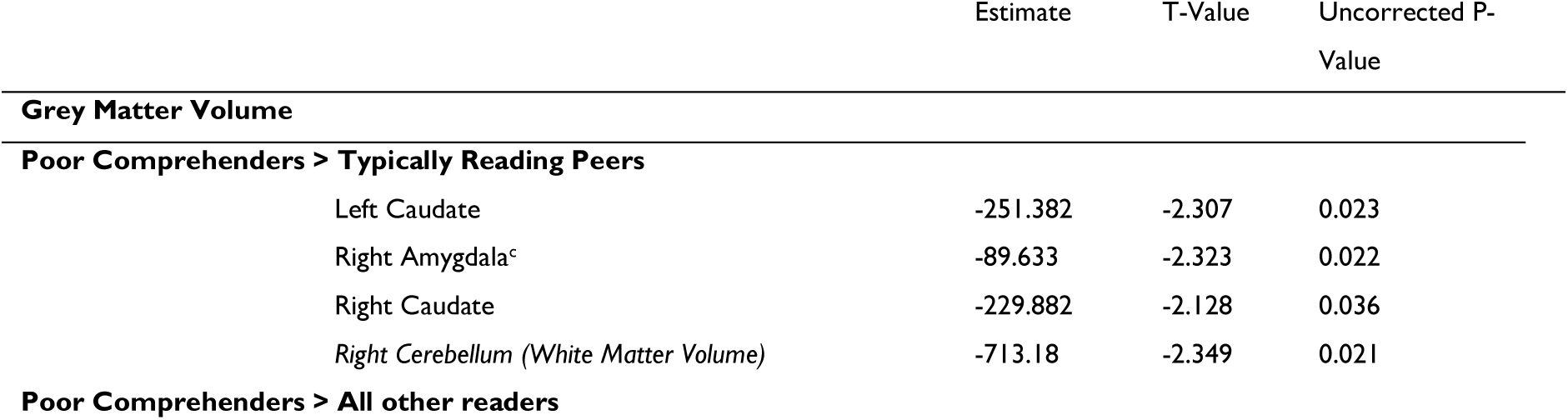
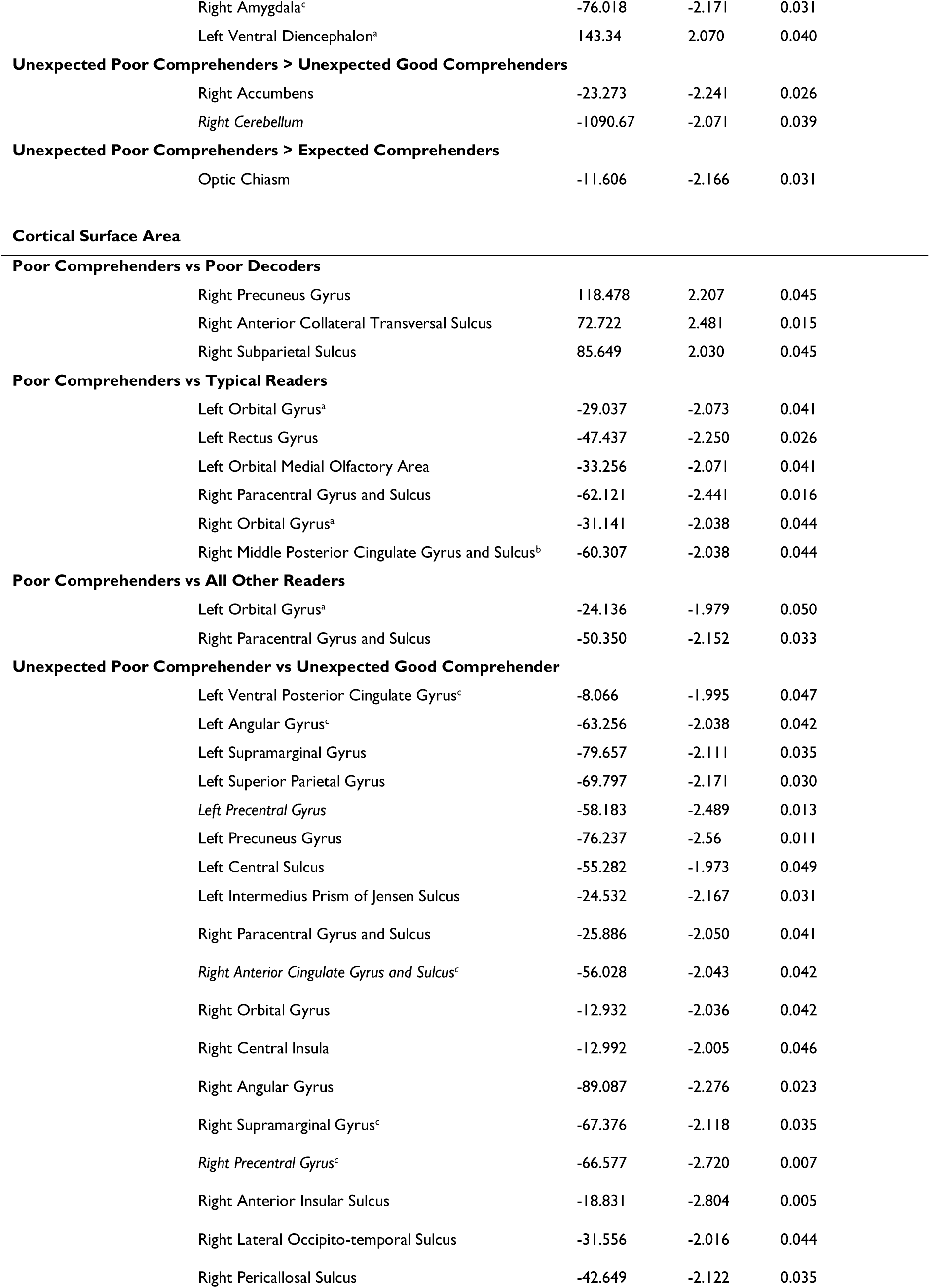
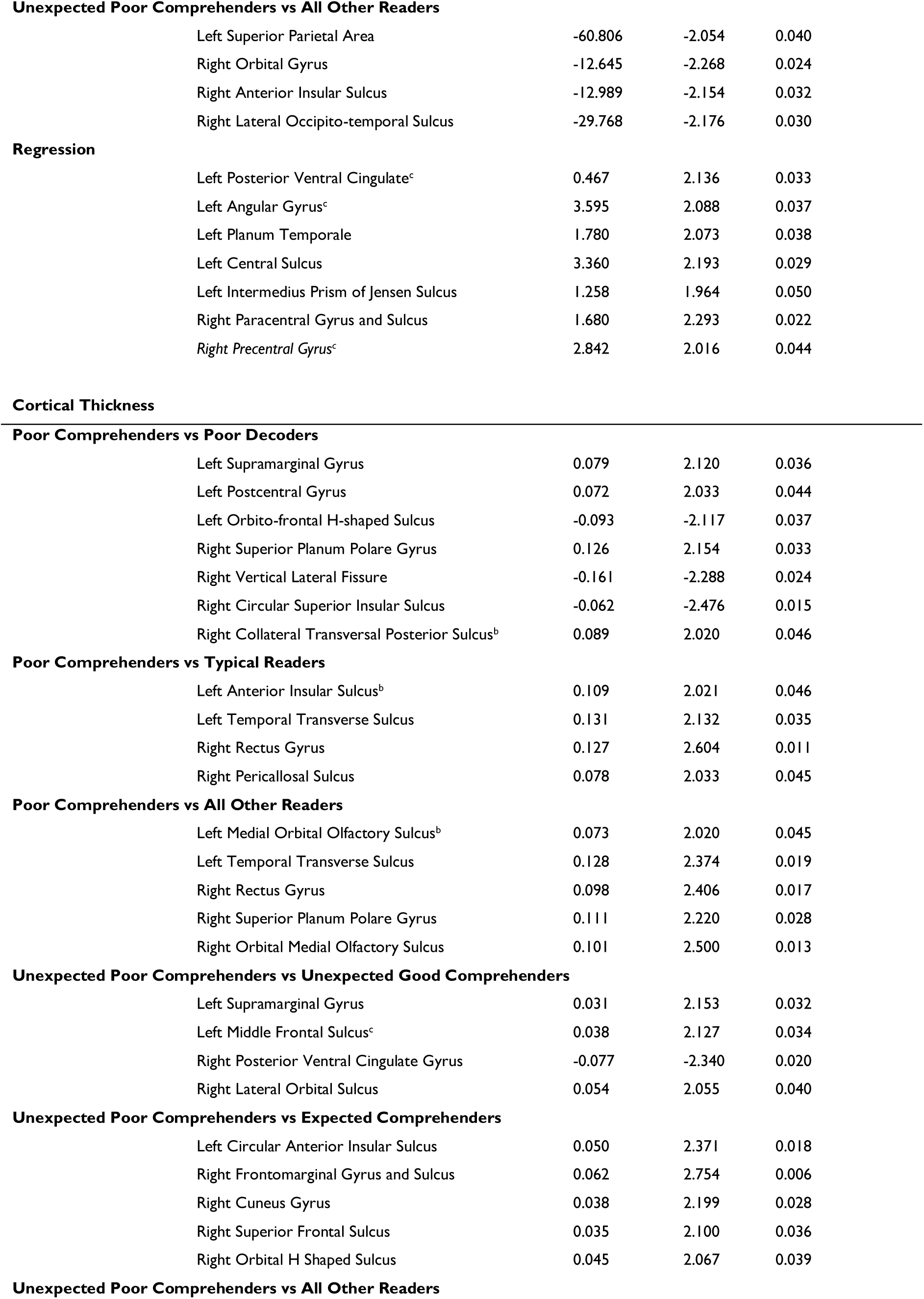
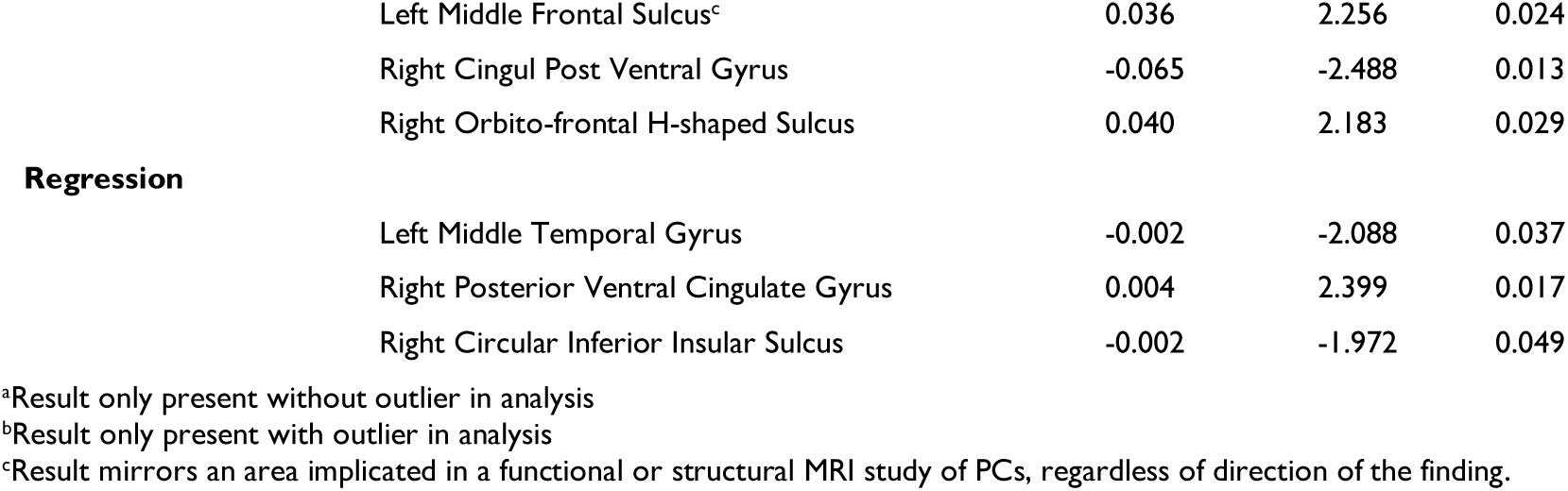
Nominally significant grey matter results

|  | Estimate | T-Value | Uncorrected P-Value |
| --- | --- | --- | --- |
| **Grey Matter Volume** |  |  |  |
| **Poor Comprehenders > Typically Reading Peers** |  |  |  |
| Left Caudate | -251.382 | -2.307 | 0.023 |
| Right Amygdala<sup>c</sup> | -89.633 | -2.323 | 0.022 |
| Right Caudate | -229.882 | -2.128 | 0.036 |
| *Right Cerebellum (White Matter Volume)* | -713.18 | -2.349 | 0.021 |
| **Poor Comprehenders > All other readers** |  |  |  |
| Right Amygdala<sup>c</sup> | -76.018 | -2.171 | 0.031 |
| Left Ventral Diencephalon<sup>a</sup> | 143.34 | 2.070 | 0.040 |
| **Unexpected Poor Comprehenders > Unexpected Good Comprehenders** |  |  |  |
| Right Accumbens | -23.273 | -2.241 | 0.026 |
| *Right Cerebellum* | -1090.67 | -2.071 | 0.039 |
| **Unexpected Poor Comprehenders > Expected Comprehenders** |  |  |  |
| Optic Chiasm | -11.606 | -2.166 | 0.031 |
| **Cortical Surface Area** |  |  |  |
| **Poor Comprehenders vs Poor Decoders** |  |  |  |
| Right Precuneus Gyrus | 118.478 | 2.207 | 0.045 |
| Right Anterior Collateral Transversal Sulcus | 72.722 | 2.481 | 0.015 |
| Right Subparietal Sulcus | 85.649 | 2.030 | 0.045 |
| **Poor Comprehenders vs Typical Readers** |  |  |  |
| Left Orbital Gyrus<sup>a</sup> | -29.037 | -2.073 | 0.041 |
| Left Rectus Gyrus | -47.437 | -2.250 | 0.026 |
| Left Orbital Medial Olfactory Area | -33.256 | -2.071 | 0.041 |
| Right Paracentral Gyrus and Sulcus | -62.121 | -2.441 | 0.016 |
| Right Orbital Gyrus<sup>a</sup> | -31.141 | -2.038 | 0.044 |
| Right Middle Posterior Cingulate Gyrus and Sulcus<sup>b</sup> | -60.307 | -2.038 | 0.044 |
| **Poor Comprehenders vs All Other Readers** |  |  |  |
| Left Orbital Gyrus<sup>a</sup> | -24.136 | -1.979 | 0.050 |
| Right Paracentral Gyrus and Sulcus | -50.350 | -2.152 | 0.033 |
| **Unexpected Poor Comprehender vs Unexpected Good Comprehender** |  |  |  |
| Left Ventral Posterior Cingulate Gyrus<sup>c</sup> | -8.066 | -1.995 | 0.047 |
| Left Angular Gyrus<sup>c</sup> | -63.256 | -2.038 | 0.042 |
| Left Supramarginal Gyrus | -79.657 | -2.111 | 0.035 |
| Left Superior Parietal Gyrus | -69.797 | -2.171 | 0.030 |
| *Left Precentral Gyrus* | -58.183 | -2.489 | 0.013 |
| Left Precuneus Gyrus | -76.237 | -2.56 | 0.011 |
| Left Central Sulcus | -55.282 | -1.973 | 0.049 |
| Left Intermedius Prism of Jensen Sulcus | -24.532 | -2.167 | 0.031 |
| Right Paracentral Gyrus and Sulcus | -25.886 | -2.050 | 0.041 |
| *Right Anterior Cingulate Gyrus and Sulcus<sup>c</sup>* | -56.028 | -2.043 | 0.042 |
| Right Orbital Gyrus | -12.932 | -2.036 | 0.042 |
| Right Central Insula | -12.992 | -2.005 | 0.046 |
| Right Angular Gyrus | -89.087 | -2.276 | 0.023 |
| Right Supramarginal Gyrus<sup>c</sup> | -67.376 | -2.118 | 0.035 |
| *Right Precentral Gyrus<sup>c</sup>* | -66.577 | -2.720 | 0.007 |
| Right Anterior Insular Sulcus | -18.831 | -2.804 | 0.005 |
| Right Lateral Occipito-temporal Sulcus | -31.556 | -2.016 | 0.044 |
| Right Pericallosal Sulcus | -42.649 | -2.122 | 0.035 |
| **Unexpected Poor Comprehenders vs All Other Readers** |  |  |  |
| Left Superior Parietal Area | -60.806 | -2.054 | 0.040 |
| Right Orbital Gyrus | -12.645 | -2.268 | 0.024 |
| Right Anterior Insular Sulcus | -12.989 | -2.154 | 0.032 |
| Right Lateral Occipito-temporal Sulcus | -29.768 | -2.176 | 0.030 |
| **Regression** |  |  |  |
| Left Posterior Ventral Cingulate<sup>c</sup> | 0.467 | 2.136 | 0.033 |
| Left Angular Gyrus<sup>c</sup> | 3.595 | 2.088 | 0.037 |
| Left Planum Temporale | 1.780 | 2.073 | 0.038 |
| Left Central Sulcus | 3.360 | 2.193 | 0.029 |
| Left Intermedius Prism of Jensen Sulcus | 1.258 | 1.964 | 0.050 |
| Right Paracentral Gyrus and Sulcus | 1.680 | 2.293 | 0.022 |
| Right Precentral Gyrus<sup>c</sup> | 2.842 | 2.016 | 0.044 |
| **Cortical Thickness****Poor Comprehenders vs Poor Decoders** |  |  |  |
| Left Supramarginal Gyrus | 0.079 | 2.120 | 0.036 |
| Left Postcentral Gyrus | 0.072 | 2.033 | 0.044 |
| Left Orbito-frontal H-shaped Sulcus | -0.093 | -2.117 | 0.037 |
| Right Superior Planum Polare Gyrus | 0.126 | 2.154 | 0.033 |
| Right Vertical Lateral Fissure | -0.161 | -2.288 | 0.024 |
| Right Circular Superior Insular Sulcus | -0.062 | -2.476 | 0.015 |
| Right Collateral Transversal Posterior Sulcus<sup>b</sup> | 0.089 | 2.020 | 0.046 |
| **Poor Comprehenders vs Typical Readers** |  |  |  |
| Left Anterior Insular Sulcus<sup>b</sup> | 0.109 | 2.021 | 0.046 |
| Left Temporal Transverse Sulcus | 0.131 | 2.132 | 0.035 |
| Right Rectus Gyrus | 0.127 | 2.604 | 0.011 |
| Right Pericallosal Sulcus | 0.078 | 2.033 | 0.045 |
| **Poor Comprehenders vs All Other Readers** |  |  |  |
| Left Medial Orbital Olfactory Sulcus<sup>b</sup> | 0.073 | 2.020 | 0.045 |
| Left Temporal Transverse Sulcus | 0.128 | 2.374 | 0.019 |
| Right Rectus Gyrus | 0.098 | 2.406 | 0.017 |
| Right Superior Planum Polare Gyrus | 0.111 | 2.220 | 0.028 |
| Right Orbital Medial Olfactory Sulcus | 0.101 | 2.500 | 0.013 |
| **Unexpected Poor Comprehenders vs Unexpected Good Comprehenders** |  |  |  |
| Left Supramarginal Gyrus | 0.031 | 2.153 | 0.032 |
| Left Middle Frontal Sulcus<sup>c</sup> | 0.038 | 2.127 | 0.034 |
| Right Posterior Ventral Cingulate Gyrus | -0.077 | -2.340 | 0.020 |
| Right Lateral Orbital Sulcus | 0.054 | 2.055 | 0.040 |
| **Unexpected Poor Comprehenders vs Expected Comprehenders** |  |  |  |
| Left Circular Anterior Insular Sulcus | 0.050 | 2.371 | 0.018 |
| Right Frontomarginal Gyrus and Sulcus | 0.062 | 2.754 | 0.006 |
| Right Cuneus Gyrus | 0.038 | 2.199 | 0.028 |
| Right Superior Frontal Sulcus | 0.035 | 2.100 | 0.036 |
| Right Orbital H Shaped Sulcus | 0.045 | 2.067 | 0.039 |

**Unexpected Poor Comprehenders vs All Other Readers**
|  |  |  |  |  |
| --- | --- | --- | --- | --- |
|  | Left Middle Frontal Sulcus<sup>c</sup> | 0.036 | 2.256 | 0.024 |
|  | Right Cingul Post Ventral Gyrus | -0.065 | -2.488 | 0.013 |
|  | Right Orbito-frontal H-shaped Sulcus | 0.040 | 2.183 | 0.029 |
| **Regression** |  |  |  |  |
|  | Left Middle Temporal Gyrus | -0.002 | -2.088 | 0.037 |
|  | Right Posterior Ventral Cingulate Gyrus | 0.004 | 2.399 | 0.017 |
|  | Right Circular Inferior Insular Sulcus | -0.002 | -1.972 | 0.049 |

<sup>a</sup>Result only present without outlier in analysis
<sup>b</sup>Result only present with outlier in analysis
<sup>c</sup>Result mirrors an area implicated in a functional or structural MRI study of PCs, regardless of direction of the finding.

### Supplementary white matter results

In the current study, a large set of white matter tracts were found to systematically differ between groups (or continuously) such that poor comprehenders had poorer white matter integrity than other readers. Specific results are reported here, with poor comprehenders having been coded as 1 in all group-based analyses, and having more negative residual scores in the regression classification models. The three segments of the Superior Longitudinal Fasciculus were averaged for all comparisons.

**Supplemental Table 4.**
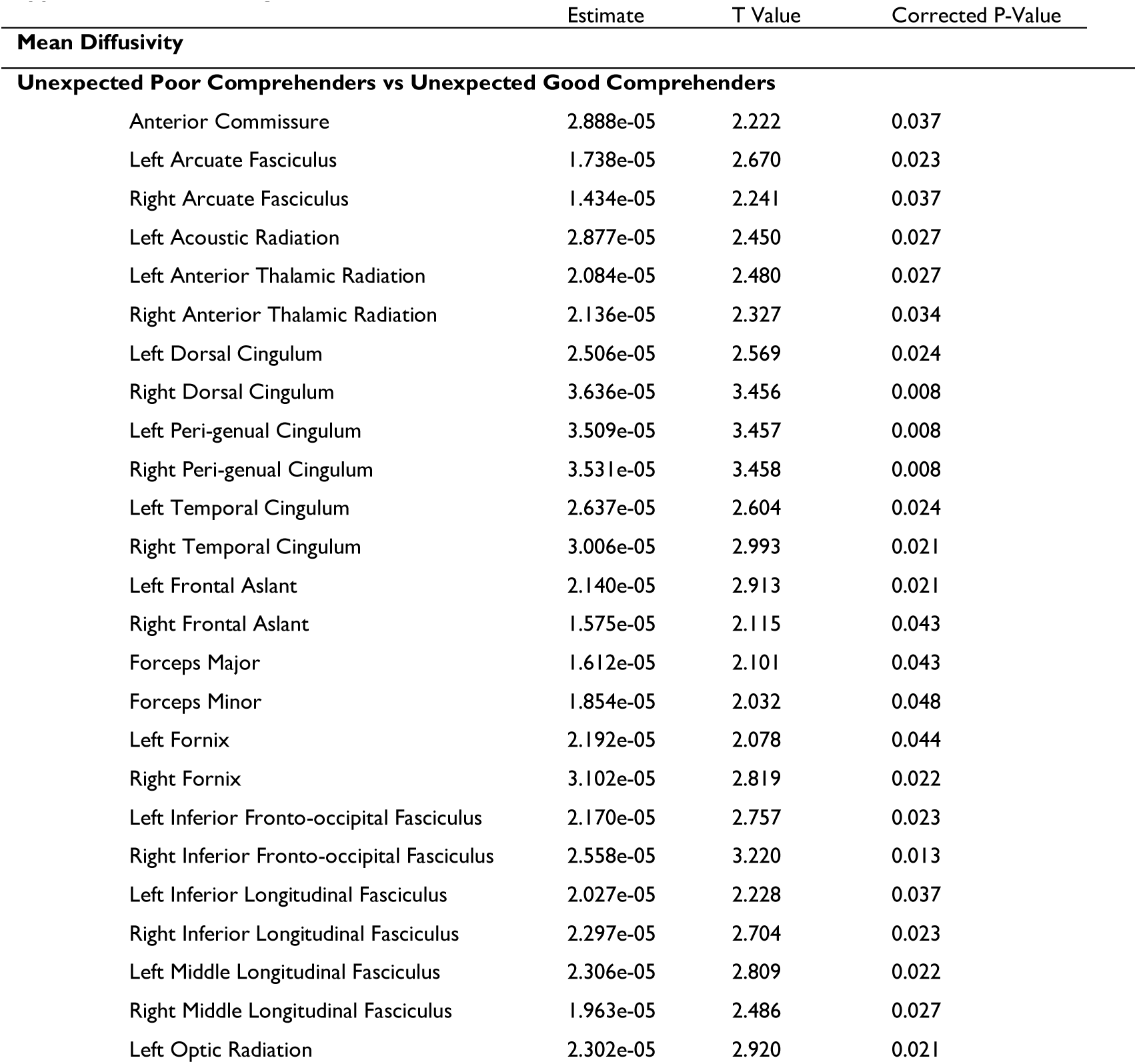
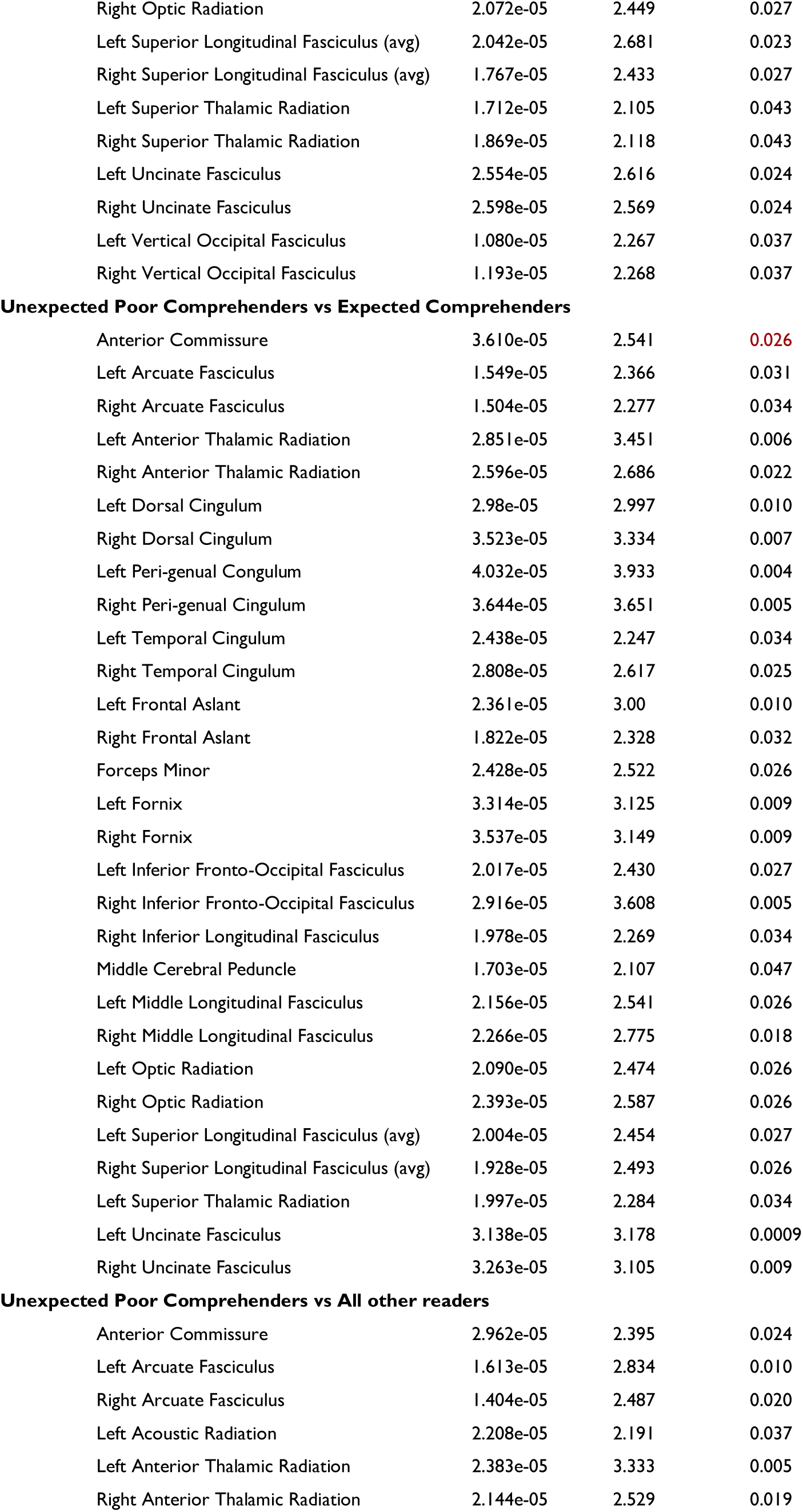
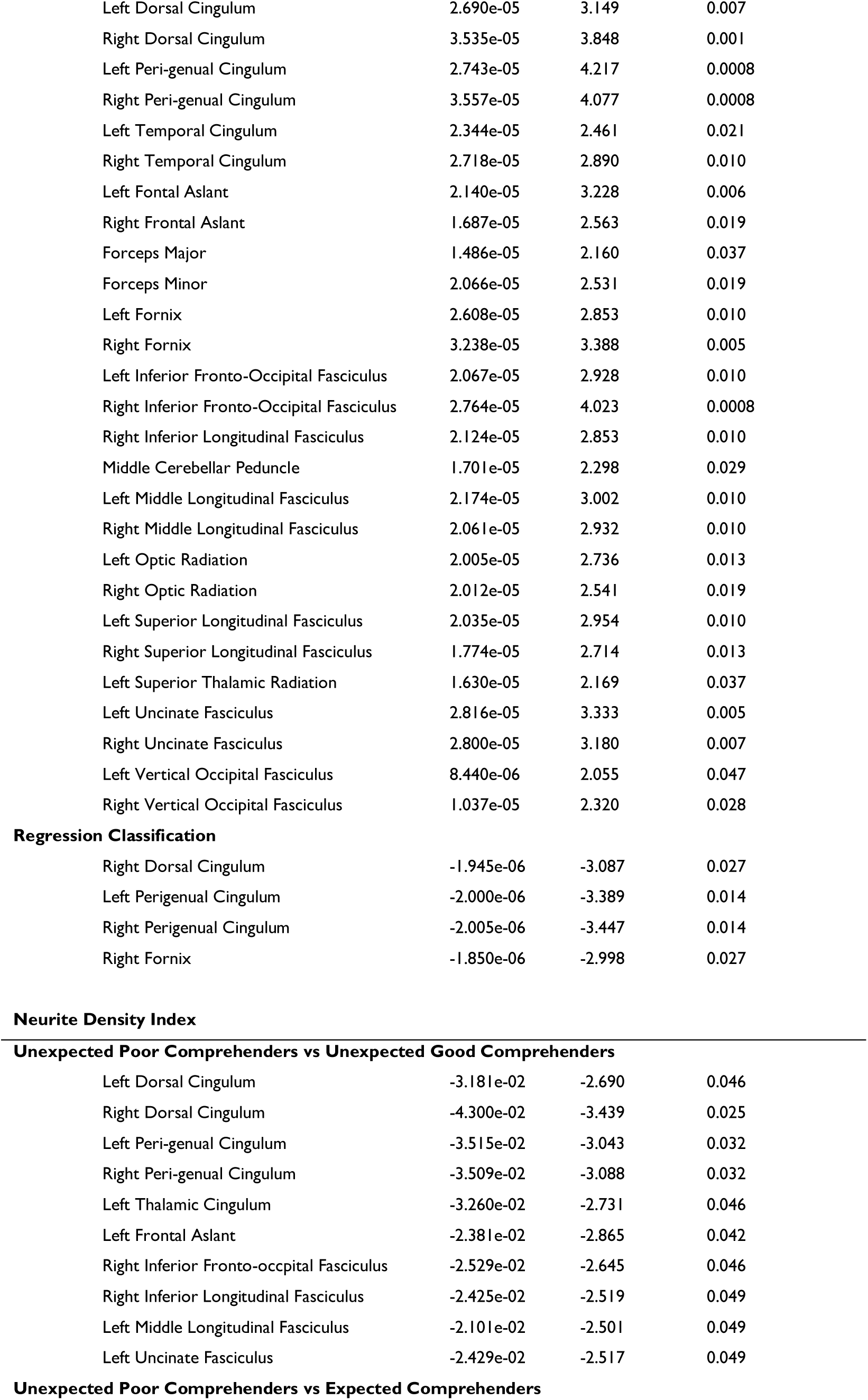
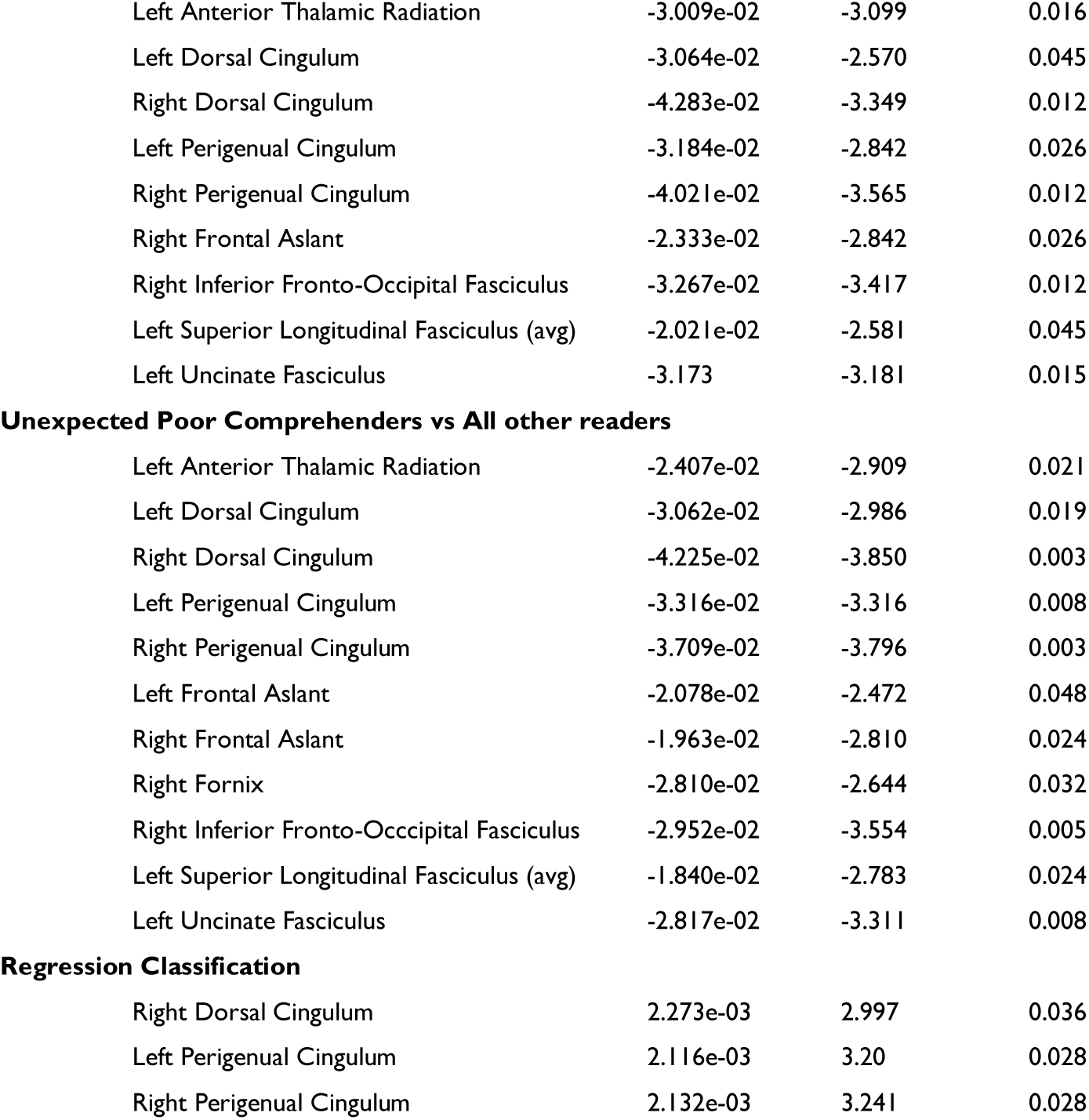
: Significant White Matter Results

|  | Estimate | T Value | Corrected P-Value |
| --- | --- | --- | --- |
| **Mean Diffusivity** |  |  |  |
| **Unexpected Poor Comprehenders vs Unexpected Good Comprehenders** |  |  |  |
| Anterior Commissure | 2.888e-05 | 2.222 | 0.037 |
| Left Arcuate Fasciculus | 1.738e-05 | 2.670 | 0.023 |
| Right Arcuate Fasciculus | 1.434e-05 | 2.241 | 0.037 |
| Left Acoustic Radiation | 2.877e-05 | 2.450 | 0.027 |
| Left Anterior Thalamic Radiation | 2.084e-05 | 2.480 | 0.027 |
| Right Anterior Thalamic Radiation | 2.136e-05 | 2.327 | 0.034 |
| Left Dorsal Cingulum | 2.506e-05 | 2.569 | 0.024 |
| Right Dorsal Cingulum | 3.636e-05 | 3.456 | 0.008 |
| Left Peri-genual Cingulum | 3.509e-05 | 3.457 | 0.008 |
| Right Peri-genual Cingulum | 3.531e-05 | 3.458 | 0.008 |
| Left Temporal Cingulum | 2.637e-05 | 2.604 | 0.024 |
| Right Temporal Cingulum | 3.006e-05 | 2.993 | 0.021 |
| Left Frontal Aslant | 2.140e-05 | 2.913 | 0.021 |
| Right Frontal Aslant | 1.575e-05 | 2.115 | 0.043 |
| Forceps Major | 1.612e-05 | 2.101 | 0.043 |
| Forceps Minor | 1.854e-05 | 2.032 | 0.048 |
| Left Fornix | 2.192e-05 | 2.078 | 0.044 |
| Right Fornix | 3.102e-05 | 2.819 | 0.022 |
| Left Inferior Fronto-occipital Fasciculus | 2.170e-05 | 2.757 | 0.023 |
| Right Inferior Fronto-occipital Fasciculus | 2.558e-05 | 3.220 | 0.013 |
| Left Inferior Longitudinal Fasciculus | 2.027e-05 | 2.228 | 0.037 |
| Right Inferior Longitudinal Fasciculus | 2.297e-05 | 2.704 | 0.023 |
| Left Middle Longitudinal Fasciculus | 2.306e-05 | 2.809 | 0.022 |
| Right Middle Longitudinal Fasciculus | 1.963e-05 | 2.486 | 0.027 |
| Left Optic Radiation | 2.302e-05 | 2.920 | 0.021 |
| Right Optic Radiation | 2.072e-05 | 2.449 | 0.027 |
| Left Superior Longitudinal Fasciculus (avg) | 2.042e-05 | 2.681 | 0.023 |
| Right Superior Longitudinal Fasciculus (avg) | 1.767e-05 | 2.433 | 0.027 |
| Left Superior Thalamic Radiation | 1.712e-05 | 2.105 | 0.043 |
| Right Superior Thalamic Radiation | 1.869e-05 | 2.118 | 0.043 |
| Left Uncinate Fasciculus | 2.554e-05 | 2.616 | 0.024 |
| Right Uncinate Fasciculus | 2.598e-05 | 2.569 | 0.024 |
| Left Vertical Occipital Fasciculus | 1.080e-05 | 2.267 | 0.037 |
| Right Vertical Occipital Fasciculus | 1.193e-05 | 2.268 | 0.037 |
| Unexpected Poor Comprehenders vs Expected Comprehenders |  |  |  |
| Anterior Commissure | 3.610e-05 | 2.541 | 0.026 |
| Left Arcuate Fasciculus | 1.549e-05 | 2.366 | 0.031 |
| Right Arcuate Fasciculus | 1.504e-05 | 2.277 | 0.034 |
| Left Anterior Thalamic Radiation | 2.851e-05 | 3.451 | 0.006 |
| Right Anterior Thalamic Radiation | 2.596e-05 | 2.686 | 0.022 |
| Left Dorsal Cingulum | 2.98e-05 | 2.997 | 0.010 |
| Right Dorsal Cingulum | 3.523e-05 | 3.334 | 0.007 |
| Left Peri-genual Congulum | 4.032e-05 | 3.933 | 0.004 |
| Right Peri-genual Cingulum | 3.644e-05 | 3.651 | 0.005 |
| Left Temporal Cingulum | 2.438e-05 | 2.247 | 0.034 |
| Right Temporal Cingulum | 2.808e-05 | 2.617 | 0.025 |
| Left Frontal Aslant | 2.361e-05 | 3.00 | 0.010 |
| Right Frontal Aslant | 1.822e-05 | 2.328 | 0.032 |
| Forceps Minor | 2.428e-05 | 2.522 | 0.026 |
| Left Fornix | 3.314e-05 | 3.125 | 0.009 |
| Right Fornix | 3.537e-05 | 3.149 | 0.009 |
| Left Inferior Fronto-Occipital Fasciculus | 2.017e-05 | 2.430 | 0.027 |
| Right Inferior Fronto-Occipital Fasciculus | 2.916e-05 | 3.608 | 0.005 |
| Right Inferior Longitudinal Fasciculus | 1.978e-05 | 2.269 | 0.034 |
| Middle Cerebral Peduncle | 1.703e-05 | 2.107 | 0.047 |
| Left Middle Longitudinal Fasciculus | 2.156e-05 | 2.541 | 0.026 |
| Right Middle Longitudinal Fasciculus | 2.266e-05 | 2.775 | 0.018 |
| Left Optic Radiation | 2.090e-05 | 2.474 | 0.026 |
| Right Optic Radiation | 2.393e-05 | 2.587 | 0.026 |
| Left Superior Longitudinal Fasciculus (avg) | 2.004e-05 | 2.454 | 0.027 |
| Right Superior Longitudinal Fasciculus (avg) | 1.928e-05 | 2.493 | 0.026 |
| Left Superior Thalamic Radiation | 1.997e-05 | 2.284 | 0.034 |
| Left Uncinate Fasciculus | 3.138e-05 | 3.178 | 0.0009 |
| Right Uncinate Fasciculus | 3.263e-05 | 3.105 | 0.009 |
| Unexpected Poor Comprehenders vs All other readers |  |  |  |
| Anterior Commissure | 2.962e-05 | 2.395 | 0.024 |
| Left Arcuate Fasciculus | 1.613e-05 | 2.834 | 0.010 |
| Right Arcuate Fasciculus | 1.404e-05 | 2.487 | 0.020 |
| Left Acoustic Radiation | 2.208e-05 | 2.191 | 0.037 |
| Left Anterior Thalamic Radiation | 2.383e-05 | 3.333 | 0.005 |
| Right Anterior Thalamic Radiation | 2.144e-05 | 2.529 | 0.019 |
| Left Dorsal Cingulum | 2.690e-05 | 3.149 | 0.007 |
| Right Dorsal Cingulum | 3.535e-05 | 3.848 | 0.001 |
| Left Peri-genual Cingulum | 2.743e-05 | 4.217 | 0.0008 |
| Right Peri-genual Cingulum | 3.557e-05 | 4.077 | 0.0008 |
| Left Temporal Cingulum | 2.344e-05 | 2.461 | 0.021 |
| Right Temporal Cingulum | 2.718e-05 | 2.890 | 0.010 |
| Left Frontal Aslant | 2.140e-05 | 3.228 | 0.006 |
| Right Frontal Aslant | 1.687e-05 | 2.563 | 0.019 |
| Forceps Major | 1.486e-05 | 2.160 | 0.037 |
| Forceps Minor | 2.066e-05 | 2.531 | 0.019 |
| Left Fornix | 2.608e-05 | 2.853 | 0.010 |
| Right Fornix | 3.238e-05 | 3.388 | 0.005 |
| Left Inferior Fronto-Occipital Fasciculus | 2.067e-05 | 2.928 | 0.010 |
| Right Inferior Fronto-Occipital Fasciculus | 2.764e-05 | 4.023 | 0.0008 |
| Right Inferior Longitudinal Fasciculus | 2.124e-05 | 2.853 | 0.010 |
| Middle Cerebellar Peduncle | 1.701e-05 | 2.298 | 0.029 |
| Left Middle Longitudinal Fasciculus | 2.174e-05 | 3.002 | 0.010 |
| Right Middle Longitudinal Fasciculus | 2.061e-05 | 2.932 | 0.010 |
| Left Optic Radiation | 2.005e-05 | 2.736 | 0.013 |
| Right Optic Radiation | 2.012e-05 | 2.541 | 0.019 |
| Left Superior Longitudinal Fasciculus | 2.035e-05 | 2.954 | 0.010 |
| Right Superior Longitudinal Fasciculus | 1.774e-05 | 2.714 | 0.013 |
| Left Superior Thalamic Radiation | 1.630e-05 | 2.169 | 0.037 |
| Left Uncinate Fasciculus | 2.816e-05 | 3.333 | 0.005 |
| Right Uncinate Fasciculus | 2.800e-05 | 3.180 | 0.007 |
| Left Vertical Occipital Fasciculus | 8.440e-06 | 2.055 | 0.047 |
| Right Vertical Occipital Fasciculus | 1.037e-05 | 2.320 | 0.028 |
| Regression Classification |  |  |  |
| Right Dorsal Cingulum | -1.945e-06 | -3.087 | 0.027 |
| Left Perigenual Cingulum | -2.000e-06 | -3.389 | 0.014 |
| Right Perigenual Cingulum | -2.005e-06 | -3.447 | 0.014 |
| Right Fornix | -1.850e-06 | -2.998 | 0.027 |
| Unexpected Poor Comprehenders vs Unexpected Good Comprehenders |  |  |  |
| Left Dorsal Cingulum | -3.181e-02 | -2.690 | 0.046 |
| Right Dorsal Cingulum | -4.300e-02 | -3.439 | 0.025 |
| Left Peri-genual Cingulum | -3.515e-02 | -3.043 | 0.032 |
| Right Peri-genual Cingulum | -3.509e-02 | -3.088 | 0.032 |
| Left Thalamic Cingulum | -3.260e-02 | -2.731 | 0.046 |
| Left Frontal Aslant | -2.381e-02 | -2.865 | 0.042 |
| Right Inferior Fronto-occipital Fasciculus | -2.529e-02 | -2.645 | 0.046 |
| Right Inferior Longitudinal Fasciculus | -2.425e-02 | -2.519 | 0.049 |
| Left Middle Longitudinal Fasciculus | -2.101e-02 | -2.501 | 0.049 |
| Left Uncinate Fasciculus | -2.429e-02 | -2.517 | 0.049 |
| Unexpected Poor Comprehenders vs Expected Comprehenders |  |  |  |
| Left Anterior Thalamic Radiation | -3.009e-02 | -3.099 | 0.016 |
| Left Dorsal Cingulum | -3.064e-02 | -2.570 | 0.045 |
| Right Dorsal Cingulum | -4.283e-02 | -3.349 | 0.012 |
| Left Perigenual Cingulum | -3.184e-02 | -2.842 | 0.026 |
| Right Perigenual Cingulum | -4.021e-02 | -3.565 | 0.012 |
| Right Frontal Aslant | -2.333e-02 | -2.842 | 0.026 |
| Right Inferior Fronto-Occipital Fasciculus | -3.267e-02 | -3.417 | 0.012 |
| Left Superior Longitudinal Fasciculus (avg) | -2.021e-02 | -2.581 | 0.045 |
| Left Uncinate Fasciculus | -3.173 | -3.181 | 0.015 |
| **Unexpected Poor Comprehenders vs All other readers** |  |  |  |
| Left Anterior Thalamic Radiation | -2.407e-02 | -2.909 | 0.021 |
| Left Dorsal Cingulum | -3.062e-02 | -2.986 | 0.019 |
| Right Dorsal Cingulum | -4.225e-02 | -3.850 | 0.003 |
| Left Perigenual Cingulum | -3.316e-02 | -3.316 | 0.008 |
| Right Perigenual Cingulum | -3.709e-02 | -3.796 | 0.003 |
| Left Frontal Aslant | -2.078e-02 | -2.472 | 0.048 |
| Right Frontal Aslant | -1.963e-02 | -2.810 | 0.024 |
| Right Fornix | -2.810e-02 | -2.644 | 0.032 |
| Right Inferior Fronto-Occipital Fasciculus | -2.952e-02 | -3.554 | 0.005 |
| Left Superior Longitudinal Fasciculus (avg) | -1.840e-02 | -2.783 | 0.024 |
| Left Uncinate Fasciculus | -2.817e-02 | -3.311 | 0.008 |
| **Regression Classification** |  |  |  |
| Right Dorsal Cingulum | 2.273e-03 | 2.997 | 0.036 |
| Left Perigenual Cingulum | 2.116e-03 | 3.20 | 0.028 |
| Right Perigenual Cingulum | 2.132e-03 | 3.241 | 0.028 |

In addition to results which were significant after multiple comparisons correction, there were some additional results which were nominally significant (significant before multiple comparisons correction and p<0.10 after multiple comparisons correction).

**Supplemental Table 5.**
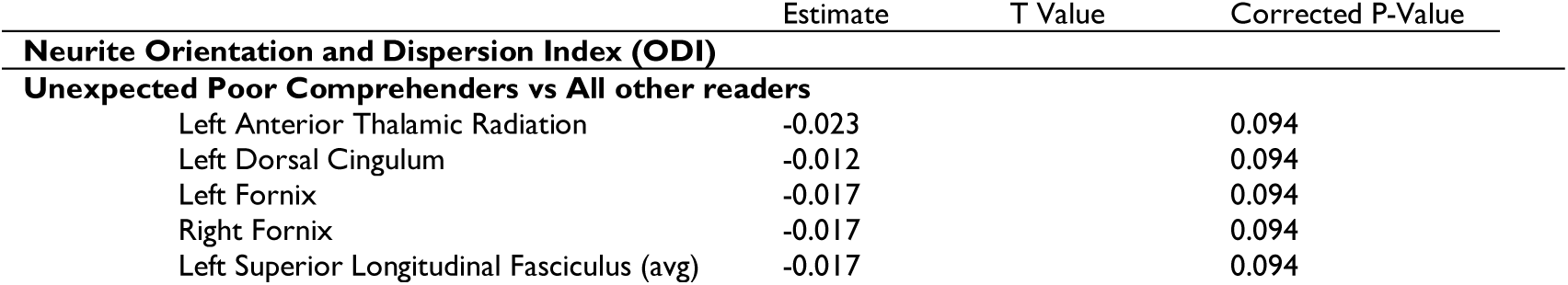
: Nominally Significant White Matter Results

|  | Estimate | T Value | Corrected P-Value |
| --- | --- | --- | --- |
| **Neurite Orientation and Dispersion Index (ODI)** |  |  |  |
| **Unexpected Poor Comprehenders vs All other readers** |  |  |  |
| Left Anterior Thalamic Radiation | -0.023 |  | 0.094 |
| Left Dorsal Cingulum | -0.012 |  | 0.094 |
| Left Fornix | -0.017 |  | 0.094 |
| Right Fornix | -0.017 |  | 0.094 |
| Left Superior Longitudinal Fasciculus (avg) | -0.017 |  | 0.094 |

### T-Tests comparing groups by classification method

One goal of the current study was to compare brain and behavior findings across three common classification methods for poor comprehenders. Groups differed in terms of their behavior, with the cutoff classification producing more distinct groups than the missed regression and regression classification methods. These t-tests compare the poor comprehender, poor decoder, and typical reader groups on decoding, oral vocabulary, and reading comprehension.

**Supplemental Table 6:**
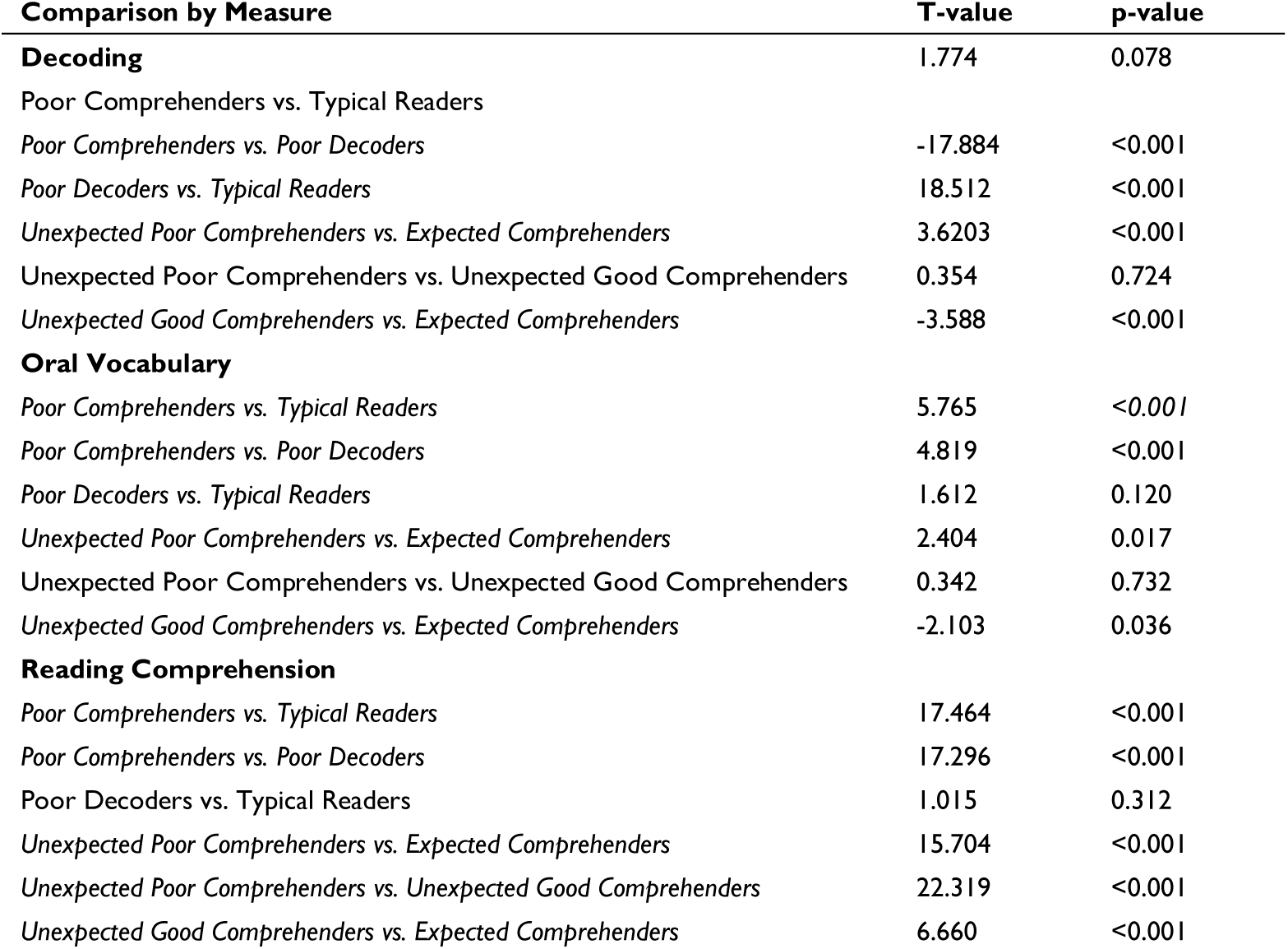
T-Tests comparing decoding, oral vocabulary, and reading comprehension across groups.

### Investigating linearity assumption of brain metric across classification approaches

Our regression classification assumed a linear relationship between residual values and morphometric measures. Only a small number of white matter tracts identified as significant in the mixed regression classification were also identified as significant predictors of white matter tractography in the regression classification. Upon closer examination of these models, it appears that less extreme morphometric measures in the unexpected good comprehender group is often driving these lack of significant results due to skewing the relationship to be non-linear across the full distribution of readers. This exemplary set of graphs illustrates this point, with readers from the mixed regression included in color (unexpected poor comprehenders in purple, expected comprehenders in turquoise, and unexpected good comprehenders in yellow) and the middle quintiles which are dropped in the mixed regression but retained in the full regression models in grey. The mean of each group is depicted in the appropriately colored cross (bar height = +/-1 SEM). Models which were found to be significantly different in both the missed regression classification and the regression classification are marked with a ^, and models which were significantly different in the mixed regression but not in the regression classification are marked with an * for ease of comparing significant and non-significant plots.

**Supplemental Figure 1.**
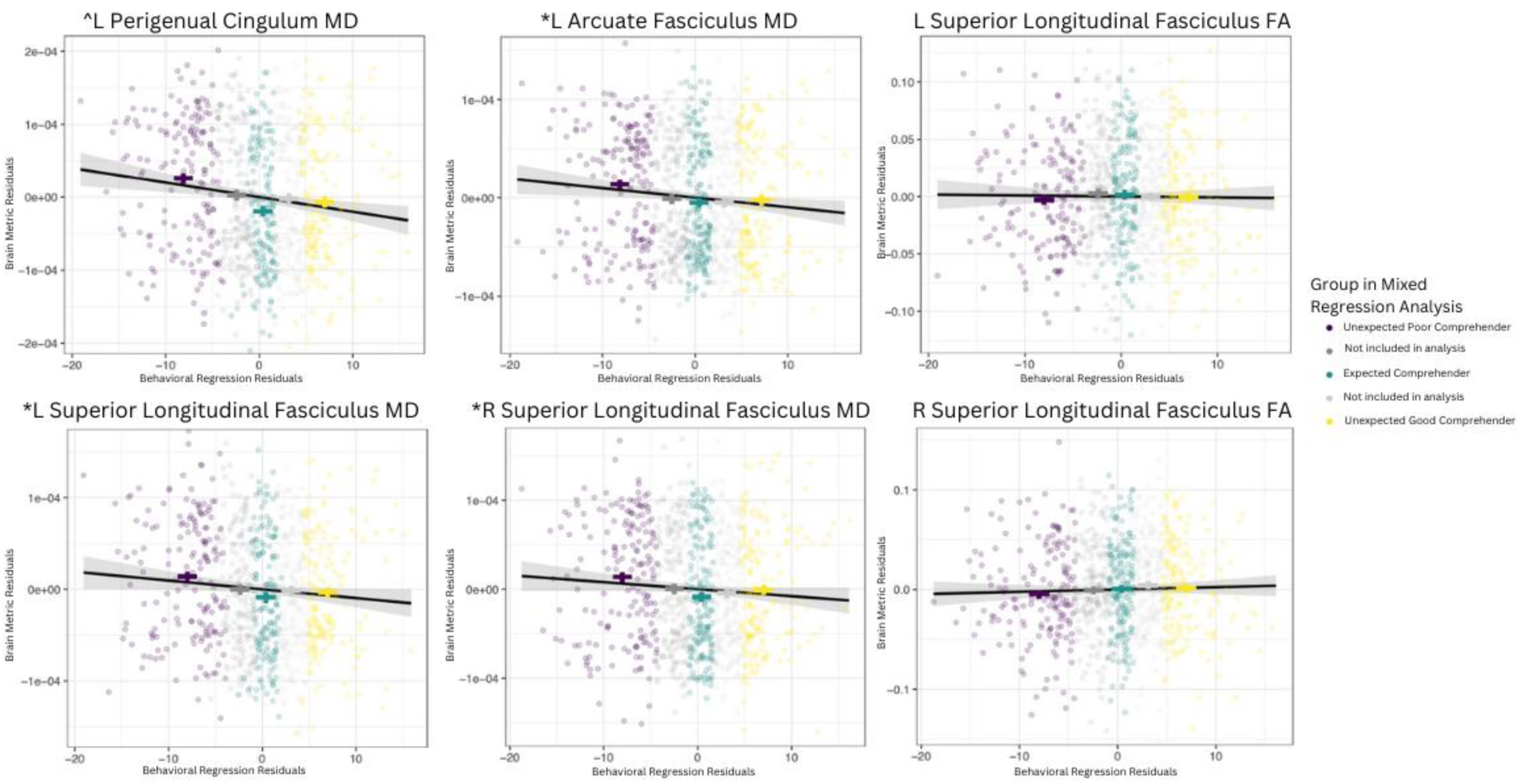
: Exemplary graphs comparing brain metrics to explore linearity across groups.

## Notes

### Competing Interest Statement

The authors have declared no competing interest.

